# Causal inference with observational data and unobserved confounding variables

**DOI:** 10.1101/2024.02.26.582072

**Authors:** Jarrett E. K. Byrnes, Laura E. Dee

## Abstract

Experiments have long been the gold standard for causal inference in Ecology. Observational data has been primarily used to validate experimental results or to find patterns that inspire experiments – not for causal inference. As ecology tackles progressively larger problems, we are moving beyond the scales at which randomized controlled experiments are feasible. Using observational data for causal inference raises the problem of confounding variables, those affecting both a causal variable and response of interest. Unmeasured confounders lead to statistical bias, creating spurious correlations and masking true causal relationships. To combat this Omitted Variable Bias, other disciplines have developed rigorous approaches for causal inference from observational data addressing the problems of confounders. We show how Ecologists can harness some of these methods: identifying confounders via causal diagrams, using nested sampling designs, and statistical designs that address omitted variable bias for causal inference. Using a motivating example of warming effects on intertidal snails, we show how current methods in Ecology (e.g., mixed models) produce incorrect inferences, and how methods presented here outperform them, reducing bias and increasing statistical power. Our goal is to enable the widespread use of observational data as tool for causal inference for the next generation of Ecological studies.

## Introduction

As Ecology advances to address problems at scales from the continental to global, we are putting our theories to the test like never before with unprecedented streams of data. With these observational data, we desire to answer questions about causal relationships to either test theory at scale or inform ecosystem management. Classically in Ecology, understanding causal relationships has been the domain of experiments. Experiments, however, have limitations for generalizing to large scales and contexts beyond study conditions. To scale up inferences to natural and managed systems, we must rapidly move beyond a scale where ideal randomized experiments are possible and responsibly seize the opportunity of new large-scale sources of observational data. Our ability to test hypotheses about causal relationships in observational data is limited, however, by two fundamental challenges: the complexity of nature and the limits of our own imaginations.

First,nature is complex! Consequently, numerous **confounding variables** – variables affecting *both* a cause and an outcome of interest (Fig. 1B and 1C) – as opposed to variables only influencing the outcome (Fig. 1A) – exist in every system. Confounding variables can lead to incorrect estimates of causal effects when not measured and controlled for in statistical analyses. Failing to control for confounding variables leads to **bias** in our statistical estimators; the estimates they yield will not be equal to their true value (Fig. 2). A simple solution for bias from confounding variables is to statistically control for confounders. Yet, this control requires knowing and measuring all the confounding variables. Even when we know which confounders to account for, collecting the data to measure and account for each and every one is likely impossible. For example, when studying plant competition, measuring all the relevant soil properties is challenged by financial or time constraints. Similarly, missing data on confounders is common with long-term survey data or in human impacted systems. Consider using historical measures of fish abundance to study the impacts of changes in biogenic habitat availability without measurements of fishing pressure during the same time-period. If fishing pressure lessened while habitat decreased, we might conclude that habitat has a negative effect on fish.

**Figure 1.**
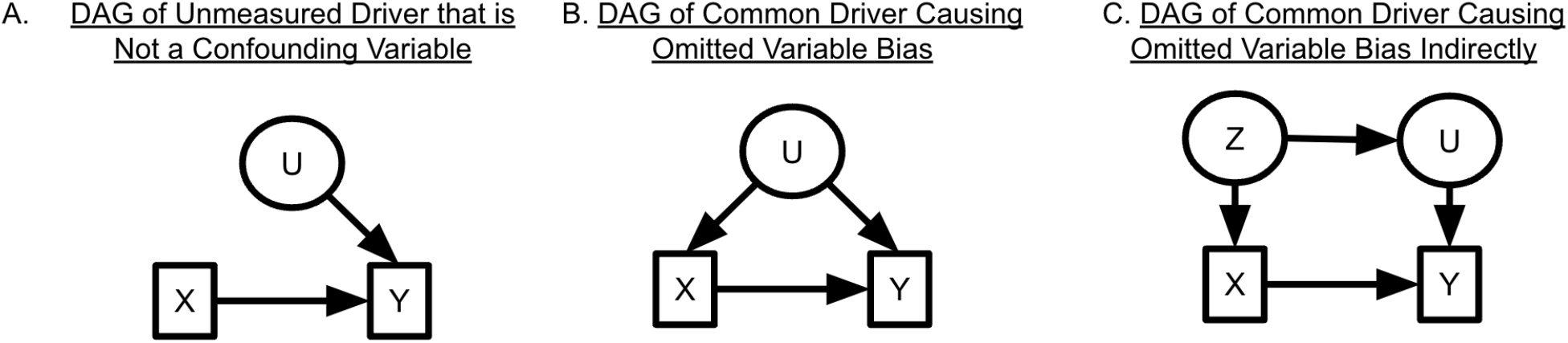
Directed Acyclic Graphs showing scenarios where unobserved variables either do not influence model results or could create problems due to confounding. A response variable of interest (*Y*) is caused by both a measured variable (*X*) and an unmeasured variable (*U*). In **(A)**, *X* and *U* are uncorrelated, and thus the lack of inclusion of *U* in a statistical model would increase the standard error of the estimate (decreases model precision) but would not lead to bias in the effect of *X* on *Y*. However, if *U* also drives *X* as in **(B)** or if *U* and *X* are driven by a common driver *Z* as in **(C)**, then omitting *U* from a statistical model causes omitted variable bias in the estimate of the effect of *X* on *Y*. Both **(B)** and **(C)** are examples of systems where the confounding common causes (*U* and *Z* respectively) must be controlled for in order to make unbiased causal inferences.

**Figure 2.**
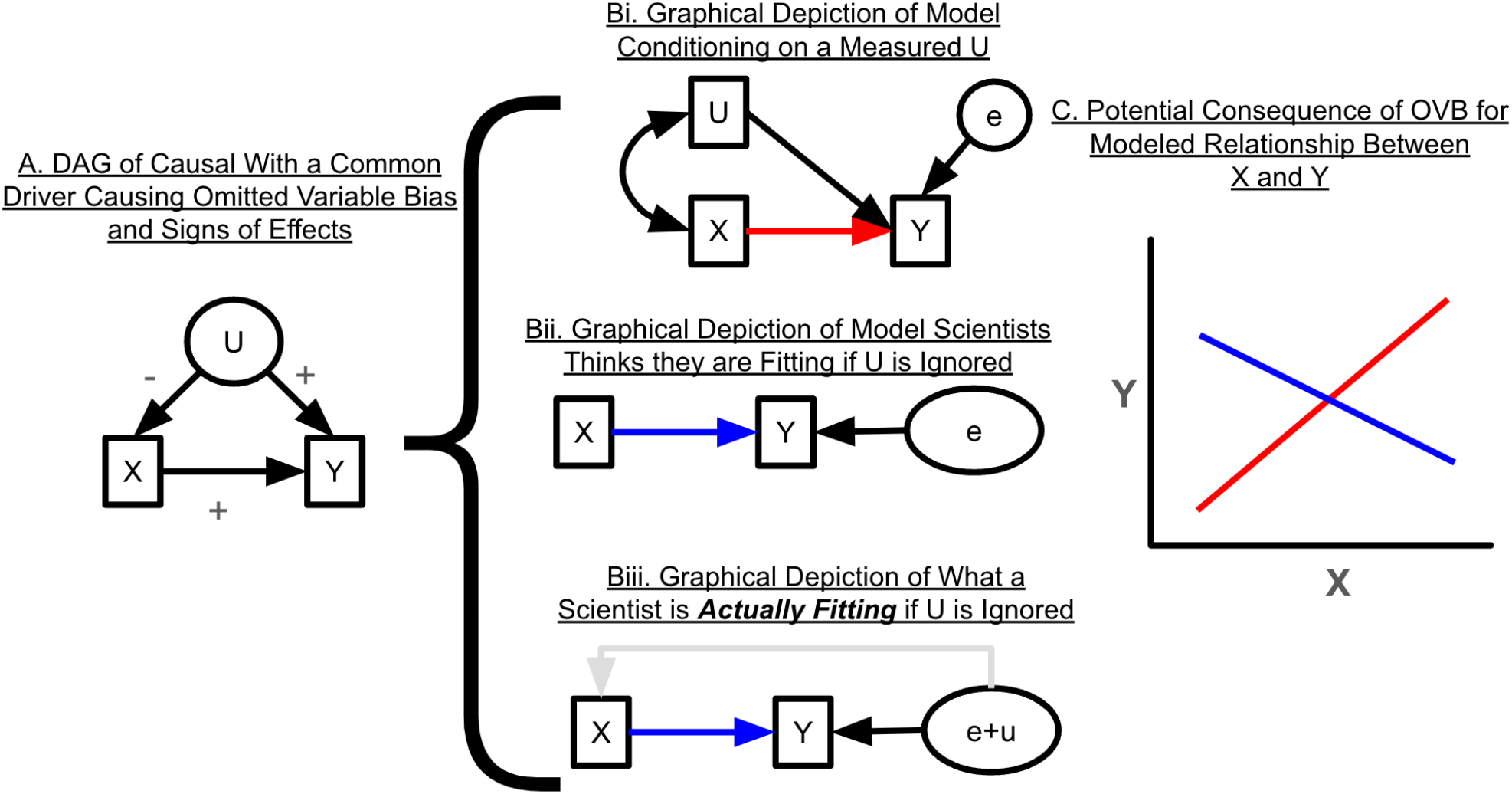
A visualization of Omitted Variable Bias and the consequences for causal inference. **(A)** shows a DAG of a system where X has a positive effect on Y, and a confounding variable *U* has a positive effect on *Y* but a negative effect on *X*. Throughout, *unobserved* (i.e., unmeasured) variables are shown in in ellipses, such as the variable *U* and the error term *e* in panel B. (**B**) illustrates different *estimations* of the DAG in **(A)** using a path analysis. See Box 1 for a brief explanation of key differences between DAGs and path diagrams. Again, we assume *U* is unmeasured. In **(Bi)**, we assume we can measure and control for U, as represented by the double-headed arrow between U and X, which represents the correlation between the two accounted for by the model. The unmeasured variable *e* is the residual sources of variation which as assumed to correlate with neither predictor. The red arrow represents the estimated path. In contrast, **(Bii)** and **(Biii)** are the reality -- where we do not have a measurement of *U* and do not control for it in the path model. The researcher thinks they are fitting the model in **(Bii)** but instead they are fitting the model in **(Biii)**, where the error term is not e alone, but rather the sum of e and variation due to the omitted variable *U*. Because of this, there is a directed path from the error term of the model to *X* (and thus *X* is endogenous). (**C)** shows the estimated relationships resulting from the models in **(Bi)** versus **(Bii)**. The lines represent the estimated relationship between *X* and *Y* from their respective models. Th red line is the true causal relationship, as estimated from **(Bi)** and the blue line contains omitted variable bias from not accounting for the confounding variable U as estimated by the model in **Bii/Biii**.

Second, as humans, we are limited in our ability to imagine how the different elements of complex ecological systems are causally related. Thinking through the entire natural history of a system to design an analysis accounting for all confounders to enable credible causal inferences from observational data is really hard, even for the most experienced researchers. As a result, causal inference from observational data is often dismissed as impossible, prompting the saying “correlation is not causation.” Dealing with the problems created by not controlling for unmeasured confounders in our statistical analyses is a first-order challenge for inferring causation from (Figs. 1, 2).

Omitting known but unmeasured, or unknown and unmeasured, confounding variables from a statistical analysis creates **omitted variable bias or OVB** (Rinella *et al*. 2020; Wooldridge 2015). **OVB** results in estimators yielding the incorrect magnitude – or even sign – of estimates (i.e., biased estimators). OVB can create spurious correlations or mask true causal relationships. OVB differs from measurement error in predictor variables, which produces a consistent bias towards zero and can be corrected for or modeled (McElreath 2020 chapter on measurement error; Schennach 2016). With OVB, we have no way of knowing the bias’s magnitude or direction without knowing all possible confounding variables and their relationships in a system. As measuring, controlling for, and even knowing all potential confounding variables is nearly impossible in complex ecological systems (*reviewed in* Dee *et al*. 2023), we are always going to miss something, threatening the validity of our causal inferences.

*Do challenges from OVB mean that we should not avoid using observational data for causal inference?* No! Rather than discounting and abandoning use of observational data for causal inference, we suggest that ecologists consider adopting well-established techniques from other disciplines that offer solutions, including psychology, economics, epidemiology, computer science, and sociology (Angrist & Pischke 2008; Heckman 2000; Hernan & Robins 2023; Holland 1986; Imbens & Rubin 2015; Morgan & Winship 2015; Pearl 2009; Robins 1989; Rubin 1974, 2005). Because these fields cannot always do experiments for logistical or ethical reasons - - for instance, it is not ethical to force a person smoke cigarettes daily to quantify the causal effect of smoking on dementia (Hernan & Robins 2023) – they have been developing tools to handle OVB for decades. Yet, these tools have been largely absent from the ecologist’s toolbox until relatively recently, with some exceptions (Arif & MacNeil 2022b, 2023; Butsic *et al*. 2017; Dee *et al*. 2023; Dudney *et al*. 2021; Grace & Irvine 2020; Larsen 2013; Larsen *et al*. 2019; MacDonald & Mordecai 2019; Rinella *et al*. 2020; Simler-Williamson & Germino 2022). If we, as a discipline, are to move to more widespread use of observational data for causal inference, we need to carefully consider the problems of OVB, the techniques we can use to mitigate them, and their assumptions.

Here, we aim to provide a guide to readily-accessible methods to cope with Omitted Variable Bias (OVB) for Ecologists. We begin describing the status quo for how ecologists most often deal with OVB. We then review tools for identifying potential sources of OVB before conducting a study or analysis, building on the foundation of using directed acyclic graphs that has become increasingly common in ecology (Arif & MacNeil 2023). To illustrate OVB and different ways to address it, we present a motivating example aiming to study the causal effect of temperature on marine snail abundances. With this example, we outline sampling and statistical designs for dealing with OVB and demonstrate them with simulations. Specifically, we compare the conclusions that would be drawn from the typical approaches an ecologist might take with data (e.g., mixed effect models, Bolker et al. 2009) to several other statistical designs that can more adequately control for omitted variables. While common approaches produce statistically biased results, our simulations demonstrate the utility of the statistical designs that are underutilized, if not novel, in ecology. We provide guidance for choosing among these designs along with a hands-on tutorial with R code for prospective users (see SI 7 for worked examples). This paper complements recent reviews in ecology of quasi-experimental methods (Arif & MacNeil 2022b; Butsic *et al*. 2017; the appendices of Dee *et al*. 2023) by expanding on cross-sectional and panel regression designs (see study design section for definitions) accounting for OVB. We hope to enable researchers to advance the field of Ecology at scale using observational data.

## How are Ecologists Coping with Omitted Variables Bias?

Confounding variables and omitted variable bias are commonly dealt with in one of five ways in Ecology. First, Ecologists use randomized controlled experiments. In an ideal randomized controlled experiment, the effect of confounding variables is eliminated when design assumptions are met. We can interpret observed effects of manipulations as causal (but see Kimmel *et al*. 2021 on why this can be difficult in practice, particularly in the field). The random assignment of treatments (e.g., Nitrogen addition) to units (plots) means that the treatment and control groups have the same level of any confounders on average. However, randomized controlled experiments are not always feasible at the scale needed to generate meaningful inferences. Further, they can impose experimental conditions that create artefacts making it difficult to generalize to nature (Ruesink 2000; Stachowicz *et al*. 2008; Wolkovich *et al*. 2012). Second, in observational studies, ecologists attempt to deal with confounding variables by measuring the confounders and controlling for them in a multivariate statistical model. Yet, as described above, measuring all confounders is often impossible – particularly in retrospective analyses of existing data where the data could have been collected previously for another purpose or question, with some or all confounders unmeasured. Moreover, all potential confounders in the system might not be known. Third, Ecologists use in mixed-effect models, which fold unmeasured cluster-level variables into random effects (Bolker *et al*. 2009; Harrison *et al*. 2018; Schielzeth & Nakagawa 2012). As we will discuss below in our section on statistical designs, if random effects are correlated with causal drivers of interest, random effects estimators are biased. Fourth, Ecologists sometimes make causal claims rooted in their knowledge of the natural history of a system. These claims can be problematic due to a lack of transparency and potential for incorrect statements about effect sizes; even the most accomplished naturalist can have gaps in their understanding of a system. Fifth, Ecologists often qualify their results verbally to avoid making causal claims – even when their research focus is causal understanding, rather than description (*but see* Laubach *et al*. 2021 on causal aims and claims). This practice muddies the waters and can create confusion over whether an author is claiming an association or implying causation while allowing themselves plausible deniability. We feel that, given our current need to understand causal relationships at large scales, these solutions are inadequate. In the worst case, they can even lead to misleading inferences. So, we turn to solutions from other disciplines to the problem of Omitted Variable Bias to augment our toolkits.

### Using Causal Diagrams to Clarify Causal Assumptions and Ferret out Omitted Variables Bias

Causal diagrams (a.k.a. Structural Causal Models from Pearl 1995; see Grace & Irvine 2020; Arif & MacNeil 2023 for introductions for Ecologists) are one of the first tools for identifying omitted variable bias (Arif & MacNeil 2023; Pearl 1995; Pearl *et al*. 2016). Causal diagrams in the form of Directed Acyclic Graphs (DAGs, see Box 1 and SI 1) visualize our understanding of causal relationships and confounding variables within a system. In doing so, DAGs transparently clarify our assumptions behind our causal claims about relationships inferred from data and show potential sources of bias from confounding variables. Critically, DAGs are assumed to include all common causes of a predictor and response of interest: including all measured and *unmeasured* confounding variables. We suggest drawing a DAGs before conducting a causal analysis – and, if possible, *before* data collection to inform which covariates might be confounding and should be measured.

#### Box 1

**An Overview of Directed Acyclic Graphs for Causal Analysis and Detecting Confounders**

Causal diagrams (e.g., Fig. 1), called Directed Acyclic Graphs (DAGs), help to determine where confounding variables might cause omitted variables bias and, in turn, to identify solutions in terms of sampling and statistical designs. In DAGs, assumed causal relationships among variables are implied by arrows with the direction of the arrows representing the direct of the causal effect. If the value of a causal variable of interest changes (e.g., via manipulation, exposure, or a natural process), there will be a concomitant change in the response variables it affects. In DAGs, these relationships are non-parametric, without functional form. Critically, for a DAG to be complete, it should include both measured and unmeasured confounding variables. We represent observed variables that can be or have been measured with boxes (e.g., *X* and *Y* in Fig. 1), and unobserved (i.e., unmeasured) variables within ellipses (*U* and *Z*).

DAGs help identify how and when to control for confounding variables. With a DAG, confounding variables can either be visually obvious or identified via software analyzing conditional independence among variables (e.g., Textor *et al*. 2016). Confounding variables can be included and controlled for directly in a statistical model or by controlling for a “child node” of that confounding variable (e.g., *U* as a child of *Z* in Fig 1C or see Fig. S1 for examples). Controlling for confounding variables helps satisfy the **back-door criterion** (Pearl 1995, Fig. S1A) for causal identification. That is, including variables that block all paths flowing from a *common cause* (e.g., *U* in 1B or Z in 1C) to both a causal variable of interest (*X*) and its response (*Y*) to “shut the back door” for causal information to flow between X and Y having nothing to do with their causal relationship. The estimate of the relationship between *X* and *Y* is then causally identified. Without controlling for confounding variables or others that block their influence in an analysis (e.g. see Fig S1), omitted confounders will cause OVB. A DAG also reveals where it is not possible to “shut the back door” due to unmeasured confounders, showing that other approaches – those presented in this manuscript, instrumental variables, Pearl’s “front door criterion,” etc. – are needed (Pearl 2009; Bellemare *et al*. 2024).

DAGs also show what variables should *<u>not</u>* be included in an analysis, such as those causing collider bias or are otherwise bad controls (e.g. conditioning on mediator variables on the causal path from X to Y). Collider bias occurs when evaluating a relationship between two variables, but conditioning on a variable they both cause. For example, conditioning on plant abundance when analyzing the relationship between disturbance intensity and herbivory intensity, but both causes plant abundance. In contrast, model selection metrics such as AIC –for predict versus causal aims - might favor including colliders or other bad controls (Arif & MacNeil 2022a). Here, we focus on bias from omitting important variables rather than including the wrong variables – which have been amply covered elsewhere (see McElreath McElreath 2020 Chapter 6; Laubach *et al*. 2021; Griffith *et al*. 2020). F

DAGs are different than path models or other graphical depictions of statistical models more common in ecology (e.g., Structural Equation Models). Both seek to show directional connections between variables but differ in several ways. 1) DAGs only represent causal relationships; path models can be causal or not. Path models can be used for non-causal aims and include unexplained correlations as double-headed arrows and other elements of error generating processes. 2) DAGs must include all common causes of the causal variable of interest and response for causal identification. 3) DAGs are non-parametric and not tied to an estimation approach; path models represent an algebraic representation of a system in the form of a statistical model. 4) Path models can include feedbacks and cycles, whereas DAGs are acyclic (see SI 2 for a discussion of feedbacks and DAGs). To show the links between DAGs and statistical models, here we present both DAGs and path models (e.g., Fig. 2).

As applied researchers, we have found that DAGs, paired with robust statistical approaches for causal inferences, clarify our own thinking and communication about ecological systems. Further, when multiple different theories suggest different DAGs, they can still be used to identify potential sources of confounding for analyses. For researchers interested in exploring possible DAGs or evaluating DAGs given some constraints, we refer readers to the field of causal discovery (Glymour *et al*. 2019).

After building a DAG as described in Box 1, one can determine potential sources of omitted variable bias, including from unmeasured confounding variables (e.g., *U* in Fig. 1B). Not controlling for confounding variables opens a “back-door” for confounding variation to flow between the causal variable and response variable of interest via an unassessed pathway (Pearl 2009). Said another way, omitting a confounding variable like *U* in Figure 1B in a statistical analysis means that it is folded into a statistical model’s error term, along with random sources of error. Figure 2 illustrates the consequences, when a confounder *<u>U</u>* has a positive effect on *X* but a negative effect on on *Y*. If we fit a model as depicted in Figure 2bi, the estimated effect of *X* on Y is positive when controlling for *U*. If we do not control for U, as in Fig. 2bii, *U* is folded into the error term, inducing a correlation between the error and *X* as seen in Fig. 2biii and leading to an incorrect estimate. Therefore, the model’s error term and causal variable of interest would be correlated, and the causal variable is **endogenous**, which is a violation of a core assumption of the Gauss-Markov theorem (Abdallah *et al*. 2015; Antonakis *et al*. 2010). This produces an incorrect estimate, shown in blue.

As an example of endogeneity and why is it problem, consider a study evaluating the relationship between nitrogen availability (*X*) and plant biomass (*Y*) across a series of fields in the context of Figure 2 as described above. Here, nitrogen availability (*X*, as in Fig. 2A) depends on field soil characteristics (*U*), and field soil characteristics also drive plant biomass (*Y*). If field soil characteristics were omitted from an analysis, then 1) the effects of soil characteristics would be included in the error term of that model so that 2) nitrogen is no longer **exogenous** (external to the system of interactions). Rather, it is **endogenous** – it is affected by elements in the error term, e.g., in a causal diagram, an arrow would go *from* error *to* nitrogen (Fig 2Biii). This endogeneity creates a correlation between the error and nitrogen in a naïve statistical model, misattributing the effect of soil characteristics to nitrogen and leading to incorrect estimates of nitrogen alone on plant biomass. We are no longer estimating the effect of nitrogen controlling for soil characteristics; therefore, the estimate of the nitrogen effect will be wrong – different from the true effect in magnitude or even sign. As discussed below, making field a random effect does not resolve this problem. If we had drawn a DAG, we could have seen where endogeneity problems like this occurred. A causal diagram is therefore a first step on the way for identifying potential OVB.

DAGs also justify choices of control variables; they make transparent the assumptions a researcher makes about how a system works for the readers of their work. They can be incorrect, however, or not include unknown confounding variables. A DAG only represents a researcher’s current understanding and own assumptions about the causal relationships within a system. Even so, they provide a useful tool to begin to think about where spatial and temporal confounders that a researcher has failed to think about might be lurking and present options to address them. Even if we do not have the correct DAG nor have thought of all possible confounders, recognizing the possibilities for spatial and temporal confounders allow us to leverage complementary approaches that lessen our reliance on a perfectly correct DAG to control for confounders. Using a case study of snails influences by temperature, let us review these approaches that combine observational sampling designs with statistical designs to control for *unobserved* and potentially *unknown* confounding variables.

### Case Study: A Problem of Omitted Snails

To illustrate these empirical challenges and potential solutions, we consider a marine benthic ecosystem modeled after the Gulf of Maine, USA, where a researcher aims to study the causal effect of temperature on snail abundance. They hypothesize that temperature influences snail metabolic and mortality rates and wish to estimate its effect on snail population abundance. Snail population abundance is also driven by recruitment, in part influenced by regional oceanography (i.e., the flow of major currents that differ in a myriad of properties). Oceanography drives both water temperature and recruitment patterns (Broitman et al. 2005; Yund et al. 2015). We assume that the researcher measured snail abundance and temperature at several sites but not recruitment or any measurement of oceanography. Thus, recruitment and oceanography are unobserved confounding variables. Estimates produced from a statistical analysis of just the temperature-snail relationship will almost certainly be wrong. Even if the researcher had measured recruitment and added it to a regression, what if there are other confounding variables lurking? Even if oceanography or recruitment were included in a statistical model, omitted variable bias remains a real possibility. The estimated effect of temperature on snails could still be incorrect: different in the magnitude or even the sign of the true effect. Fortunately, our researcher drew a DAG (Fig. 3A) and recognized that temperature at the scale of a replicate was also influenced by local variation (e.g., from the many possible sources of microclimatic variability). With this DAG in hand, they realized they could control for both observed and unobserved confounding variables with appropriate sampling and statistical designs.

**Figure 3.**
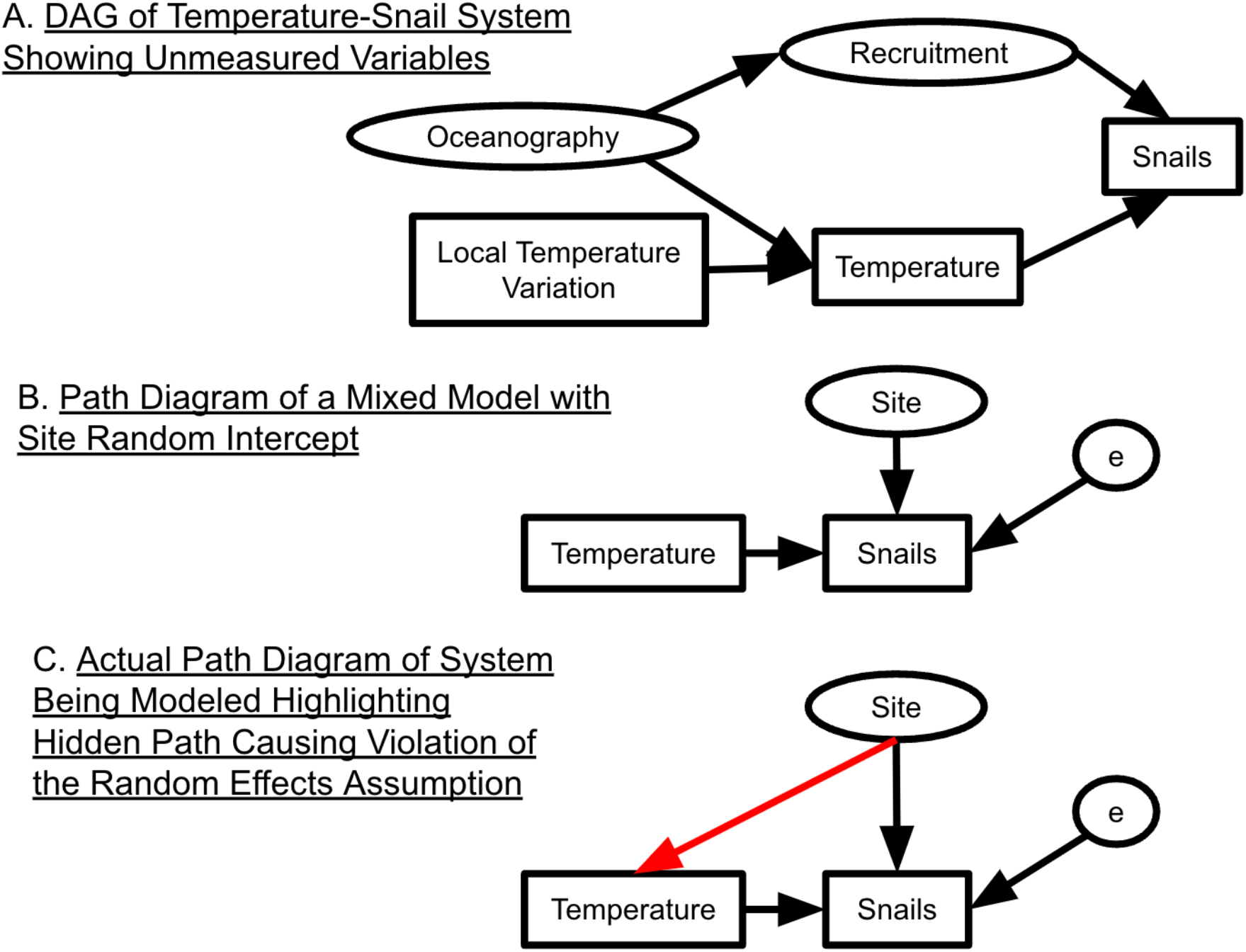
A causal diagram describing the controls of snail abundance in an intertidal ecosystem and path models of a random effects models ignoring the random effects assumption. In the system in **(A)**, oceanography drives both temperature and recruitment, and both drive snail abundance. Temperature, however, is also driven by local influences as well. This could be variability in plot-level temperature within a site – i.e., sources of variation in microclimate - or site-level temperature variability over space or time uncorrelated with local oceanography, recruitment, or other site- or plot-level confounders. The mixed model with a random effect for site in **(B)** assumes that there are no site-level drivers of temperature and does not account for the relationship between site and temperature. Thus, the effect of temperature on snails is confounded by any correlated site-level drivers that correlate with temperature at the site level. The assumption of no relationship between site and temperature contrasts with the violation of endogeneity highlighted in **(C)** showing that temperature is indeed at least partially drived by site. When we fit a mixed model, the red path in (**C**) is not included, creating an endogeneity problem and violating the Random Effects assumption. As these error variables are unobserved, we include them in an ellipse as with other unobserved variables.

### Sampling Designs that Enable Statistical Methods to Cope with Omitted Variable Bias

Multiple sampling designs for data collection enable the use of statistical designs that can address omitted variable bias from confounding variables that vary across space, time, or both. A key feature in these sampling designs is that there is some **hierarchical** or **clustered** structure to the data. The nesting of multiple observations within a cluster or group (e.g. site) can allow the causal variable of interest to vary across replicates within a cluster while the confounder varies between clusters at the cluster level (Fig. 4). This within/between partitioning enables us to control for confounding variation. Clustered data is often also referred to as a hierarchical or nested sampling design (Gelman & Hill 2006). We use these terms interchangeably.

**Figure 4.**
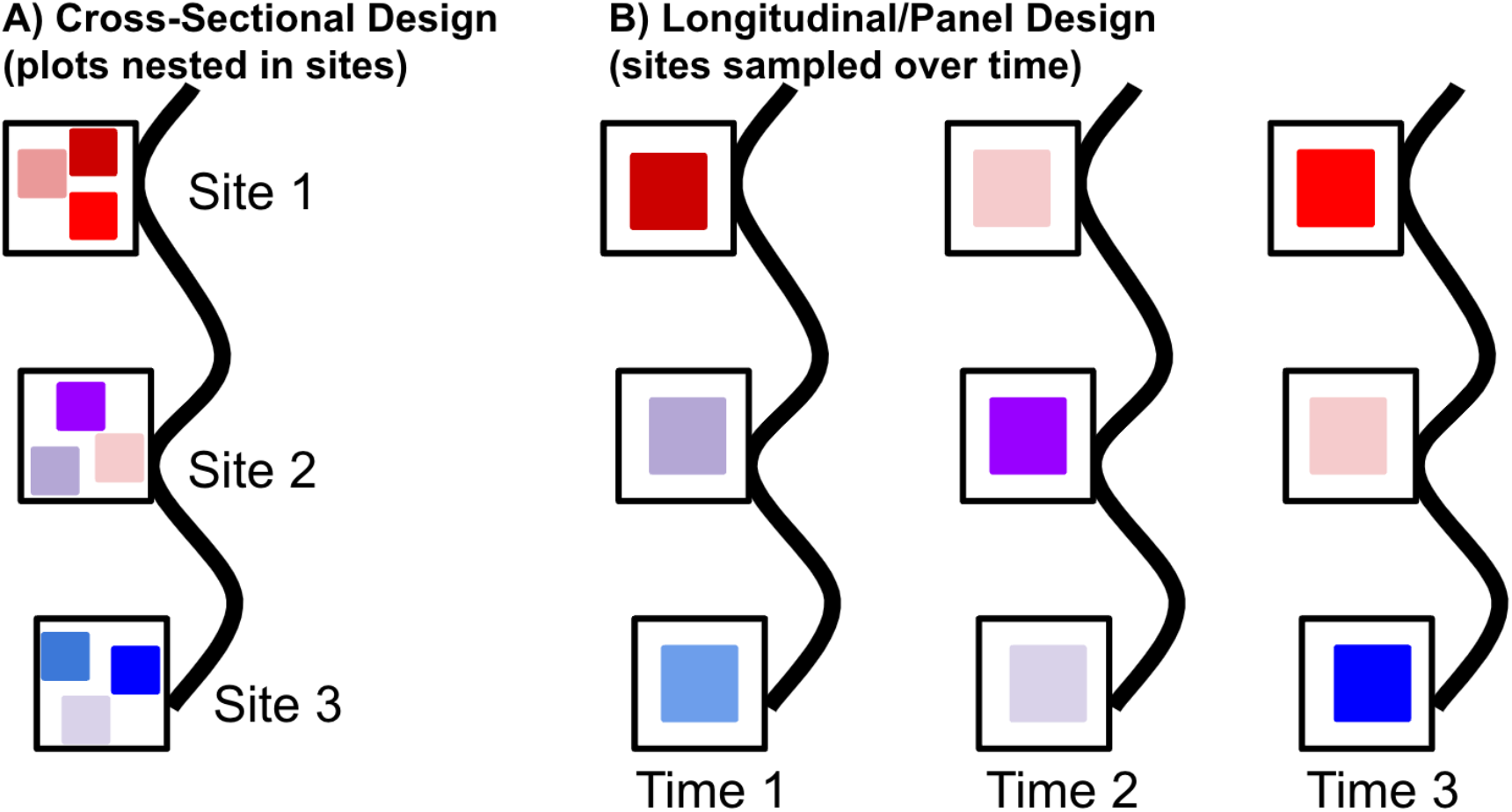
Visual examples of hierarchical study designs with plots nested within sites sampled at one point in time in A and through time in B. This figure shows sites distributed along a coastline with a corresponding thermal gradient, with one or more plots sampled within each site, depending on the design. Open squares are sites. Closed squares are plots within sites. Color of square is proportional to temperature, with red and blue signifying warm and cold respectively and other colors placed on a gradient between them. These sampling designs therefore have variation across space, as in the cross-sectional sampling design in A, or in both space and time as in B. which shows longitudinal or panel data, where the same plots within sites are observed through time. The sampling design in (A) can allow researchers to study temperature variation within sites as well as between sites. The design in (B) enables a researcher to leverage variation in space and time, including examining variation within sites through time.

Clustered sampling designs can take several forms and generate different types of variation to study. First, a sampling design could include multiple plots sampled within sites at a single point in time (Fig. 4A) – a **cross-sectional design**. When sites span environmental gradients with variation in a causal variable of interest (e.g., temperature differences), confounding variables also vary across these spatial gradients. In our case study, a spatial gradient in temperature across sites also reflects the spatial gradient in oceanography that affects both temperature and recruitment, thus confounding the causal relationship of interest between temperature and snails across sites. However, with data collected from a cross-sectional sampling design with plots within sites, we can use variation in plot temperature *within*-sites to isolate its effect on snails rather than the variation *between* sites, which contains sources of confounding variation (e.g., rockpools of different size that warm to different degrees during low tide for within versus site-level oceanographic features that drive temperature and recruitment for between).

Second, one could sample the same plots (or sites) repeatedly through time (Fig. 4B) in a **longitudinal** or **panel data design**. This sampling design enables using approaches that can leverage variation *within-sites through time*. Longitudinal data can enable many approaches to remove the effects of confounding variables that vary across sites, switching the variation we study to variation within-site (or plot) through time as opposed to between sites. Developing an understanding of how cross-sectional and panel data structures, along with variations and extensions (Box 2 and SI 3), can be used in conjunction with statistical designs to remove variation from confounders is key to confronting OVB.

### Statistical Designs to Cope with Omitted Confounders

With hierarchical or clustered (hereafter clustered) data and a DAG in hand, we turn to well-established statistical designs to handle omitted confounders for causal analysis. We emphasize the term *‘designs*’ over *‘methods’* because one could implement these statistical designs using a variety of estimation approaches (e.g., linear regression, Generalized Linear Models, as a part of Structural Equation Models, or with Bayesian techniques). These statistical designs have different costs and benefits and differ in the assumptions required for interpreting an estimate as a causal effect (also see Table S2). Yet, most of the following designs – with the exception of random effects models as shown below – allow us to flexibly control for both known and unknown confounding variables (see Angrist & Pischke 2008; Dudney *et al*. 2021; Ferraro & Miranda 2017). They are also straightforward to implement (as demonstrated in SI 7). Thus, we believe these statistical designs are a key advance worth consideration by Ecologists.

We illustrate how the different designs work using a common set of terms for causal variables of interest *(x*; e.g. local temperature), the outcome or response variable of interest (*y*; e.g. snail counts), and confounding variables (*w*; e.g. recruitment) in a regression. Our snail example includes data from different sites (*i*) sampled either at multiple time points in panel data design or in multiple plots (*j*) in the case of a cross-sectional data design as above. For the sake of simplicity, we assume a linear model form with normally distributed error (ϵ), although the framework also applies for generalized linear models. The model takes the form of

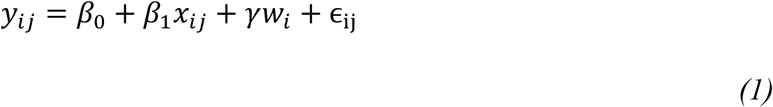

where *y*_*ij*_ is the abundance of snails at site *i* in year or plot *j*, β_0_ is the intercept – the abundance of snails if the temperature and recruitment were 0, β_1_ is the effect of temperature *x*_*ij*_ at site *i* in year or plot *j* on snails, *γ*is the effect of recruitment *w*_*i*_ at site *i* on snail abundance, and ϵ_ji_ is other drivers of snail abundance. *Thus, we have a confounding variable that varies spatially, across sites, and is correlated with average site temperature*. Our goal is to estimate *β*_1_(the effect of temperature on snail abundance) and eliminate the effect of confounding variables (and resulting bias). Due to shared oceanographic influences, *x*_*ij*_ and *w*_*i*_ are correlated. If we had measured *w*_*i*_, we could include it in our model, and by conditioning on observables with *γ*as the effect of *w* on *y*, produce a causally identified estimate of *β*_1_assuming no other confounders. Supplementary Table 1 explains coefficient definitions for all models.

Without measuring and including the confounder, *w*, as in the design above, we would instead fit the following naïve regression design with only the abundance of snails (*y*_*ij*_) as a function of temperature (*x*_*ij*_) and random error:

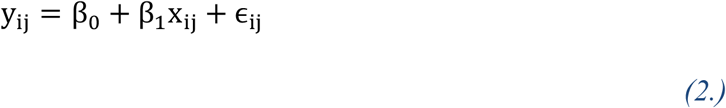

Due to omitted variable bias, our causal inference about *β*_1_would be incorrect: different from the true causal effect, because temperature (*x*_*ij*_) is endogenous – correlated to the error term – when *w*_*i*_ is included in the error term. This endogeneity problem violates the assumptions of the Gauss-Markov theorem and its extensions (Wooldridge 2015) and underlies OVB.

### What Ecologists Typically Do: Random or Mixed Effects Models That Fail to Solve OVB

Mixed effects models have been popular in ecology for the past two decades (for useful reviews, see Bolker et al. 2009, Schielzeth and Nakagawa 2012, Harrison et al. 2018). Originally used to partition variation in heritability between different relatives (Fisher 1919), **random effects –** the effects of clusters in data assumed to come from a random distribution (but see Gelman & Hill 2006 on the linguistic ambiguities surrounding fixed and random effects) – quickly became a mainstay in the partitioning of variation in randomized experiments with subsamples taken within clusters (Cochran 1937; Eisenhart 1947). They have become a standard part of the toolbox for analyzing ecological experiments (Schielzeth & Nakagawa 2012) and are frequently used when analyzing observational data in ecology.

Random effects account for clustering in data via the error structure of the model (Bolker *et al*. 2009; Gelman & Hill 2006), rather than estimating cluster means as part of the data generating process of a model (i.e., via fixed effect for each cluster’s mean, using the terminology of the mixed models literature). This results in gains in efficiency (i.e., costing fewer degrees of freedom). Further, as random effects are assumed to be drawn from a common distribution, they have benefits for analyses of unbalanced samples and for regularizing cluster means (i.e., shrinkage, drawing them towards the grand mean, see Efron & Morris 1975).

For these reasons, Ecologists conducting a study akin to our snail-temperature example would likely gravitate towards a mixed effects model to account for variation between sites in snail abundances, using a mixed effects model design such as:

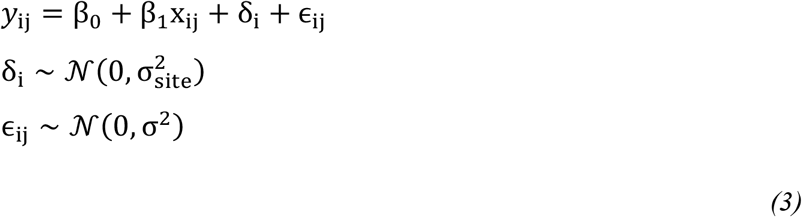

All coefficients are as in eqn. 2, with the addition of δ_i_, the random effect – a site-specific deviation of site *i* from our common intercept, β_0_, due to variation assumed to follow a normal distribution with a mean of zero and variance of 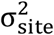. As we will see, because this is a random effect, if site is correlated with temperature, we cannot resolve the problem of OVB with this design.

### What Assumptions is a Random Effects Design Making when it Comes to Omitted Variable Bias?

Why does the above model not control for omitted confounders via its site effect? Why do random effects designs produce incorrect statistically biased results in the face of omitted confounders? To understand this problem, remember that, when we model random effects, we are not modeling group means *per se* (Robinson 1991). Rather, we are modeling correlation in our error structure due to clustering in our data (Bolker *et al*. 2009; Schielzeth & Nakagawa 2012; Wooldridge 2010). This difference – modeling error instead of modeling means *per se* – results in many of the above benefits, but also introduces one new assumption not often considered – a variation on the assumption of endogeneity we call the **Random Effects Assumption**. This assumption states that the random effects, which are part of the error term, are not correlated with any covariates in the regression (Antonakis *et al*. 2021; Wooldridge 2010).

Thus, in our snail and temperature example, in a mixed model in equation 3, the random effects of site are part of the error term and assumed to be uncorrelated with temperature for the random effects estimator to be unbiased (Schielzeth & Nakagawa 2012; Wooldridge 2010). Given the causal DAG of the system, we know this assumption is false. Therefore, in equation 3, while site is incorporated into δ_i_, the effect of temperature on snails is not causally identified. This estimator is biased due to the violation of the Random Effects Assumption; in short, estimates of β_1_will be wrong (and it is hard to know how wrong).

We can see more clearly how a mixed effects model would violate the Random Effects Assumption in Figure 3. Figure 3B shows a path diagram for a random effects model. Site effects here are site-level residuals drawn from a normal distribution. They represent all other abiotic and biotic forces happening at the site level, but they also are assumed to be uncorrelated with temperature at the site level. Given the information in Figure 3A, however, we know that this is not accurate. Therefore, the key assumption for an unbiased estimator is violated. If we were to take a step back and think about our analysis goals and our causal understanding, again representing unmeasured quantities in ellipses, what we are modeling is Figure 3C. Here, while a random site effect would yield all the benefits discussed above, we can see that site-level confounders create a violation of the Random Effects Assumption; said another way, our site-level random effect is not endogenous, and thus the random effects estimator is bias (see simulations below and Fig. 6 to see the biased estimates a random-effects model produces). We posit that violations of the Random Effects Assumption is likely common in Ecology – particularly in observational data that spans environmental gradients. The frequency and consequences of violating this assumption for Ecology is neither well explored nor acknowledged widely enough. We need solutions that do not produce biased results due to easily violated assumptions.

### Enter the Econometric Fixed Effects Design

The Econometric Fixed Effects Design represents a familiar starting point for many ecologists who are used to using categorical variables in ANOVA and ANCOVA (e.g., Gotelli & Ellison 2012). Unfortunately, there are many uses of the term “fixed effect,” leading to a wealth of confusion across fields (Gelman & Hill 2006). Here we use the term “fixed effect” as drawn from the econometrics literature, where it refers to attributes of a system (e.g., site, plot, or year) that vary by cluster (i.e., a within cluster intercept) that are encoded in models as dummy or categorical variables (e.g. representing sites or other descriptor of how our data is clustered). We also use “fixed effect” in the language of the mixed model literature – i.e., that the cluster means are estimated as part of the data generating process of the model, not as part of the random error component.

Recognizing that confounding variables vary at the cluster-level (e.g. site), we have three options to flexibly control for the effects of confounding variables. First, we can use a bit of algebra known as the **within transformation** or **fixed effects estimator** (Bell *et al*. 2018; Wooldridge 2010) that is similar to within-subjects centering in Ecology (van de Pol & Wright 2009). To illustrate, we manipulate the following equation:

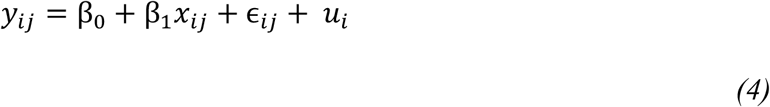

following notation of eqn. 1 where *x*_i*j*_ is our casual variable of interest, but the error term is composed of idiosyncratic (random error), ϵ_*ij*_, and *u*_*i*_, which represent differences across sites *i* including unmeasured confounding variables. As site-level confounders don’t vary across within-site replicates, 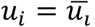. We can use this to our advantage via the **fixed effects transformation**. For this transformation, we remove the effect of site-level confounding drivers, by subtracting the site-level average value of 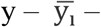 from both sides of the equation. On the right-hand side, we can average over all terms at the site level to subtract 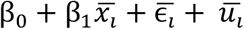 which leads to a transformed model.

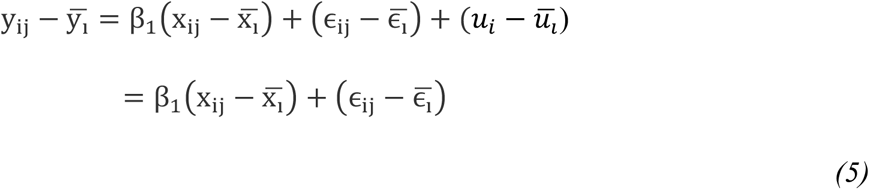

We have algebraically removed the confounding influence of time invariant site-level confounding variables contained in *u*_*i*_ – whether they were observed or not.

Second, to achieve the same effect as this fixed effects transformation (see Fig. 5A for a path diagram of the model), we could use a dummy variable for each cluster (i.e., a 0/1 encoding of *x*_*2i*_ for each cluster, an econometric fixed effect) multiplied by the cluster mean, *λ*_*i*_, or simplify the equation to just use cluster means, *λ*_*i*_, as categorical variable per site as in Fig. 5B. This is merely two ways of writing the same model.

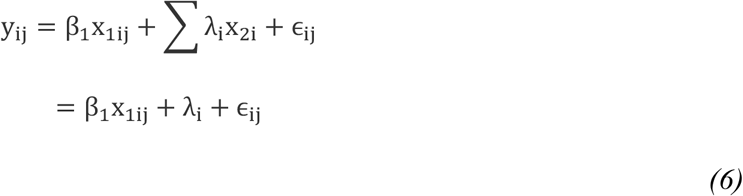

**Figure 5.**
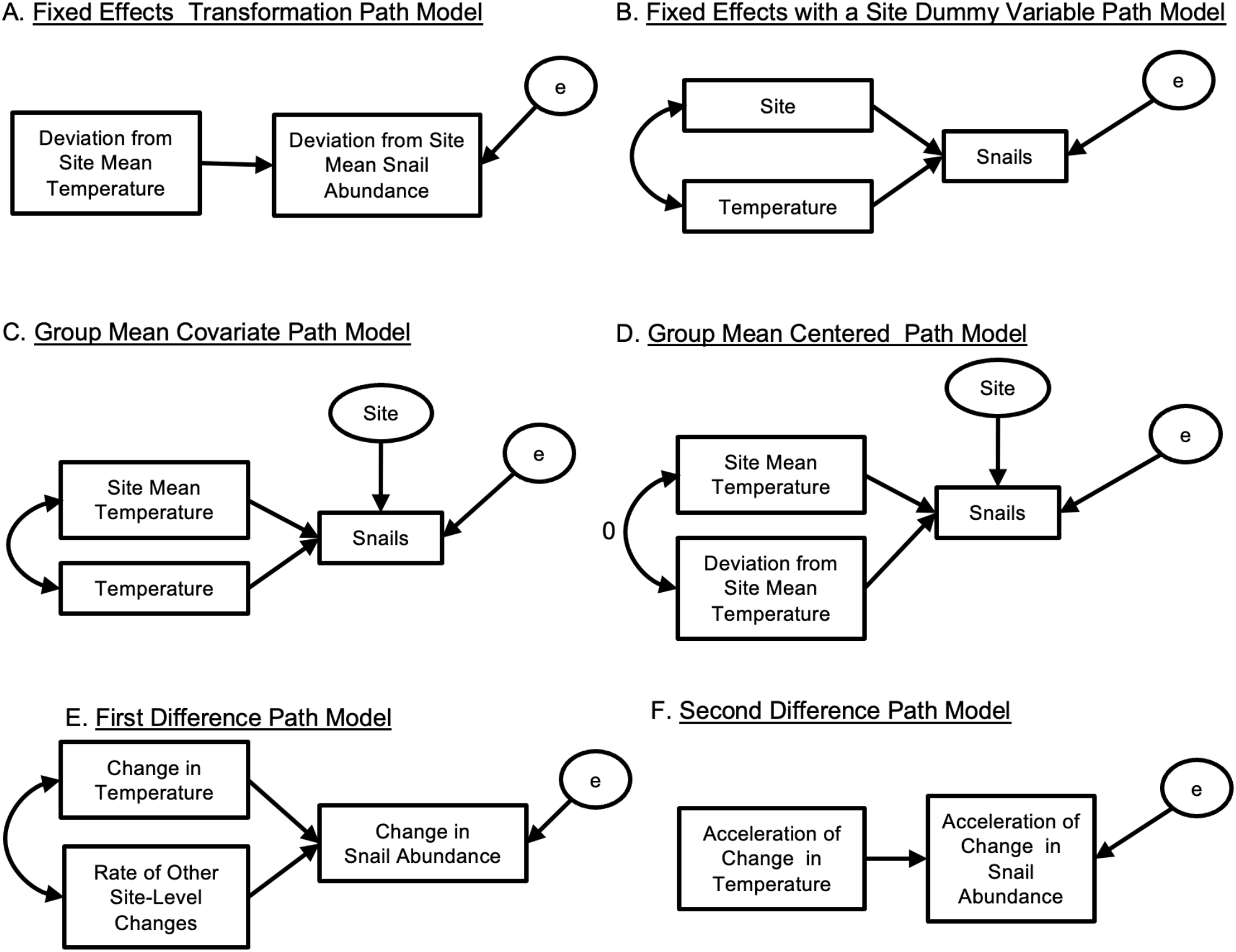
Path diagrams of different statistical models handling omitted variables in the text. (**A**) and (**B**) show two variations on the econometric fixed effect model design corresponding to equations [5] and [6] respectively. (**C**) represents the group mean covariate design in equation [7] and (**D**) represents the group mean centered design from equation [8]. Finally, **(E)** shows the first differencing approach from equation [9] and **(F)** the second differencing approach. As the true error variables are unobserved, we include them in an ellipse.

**Figure 6.**
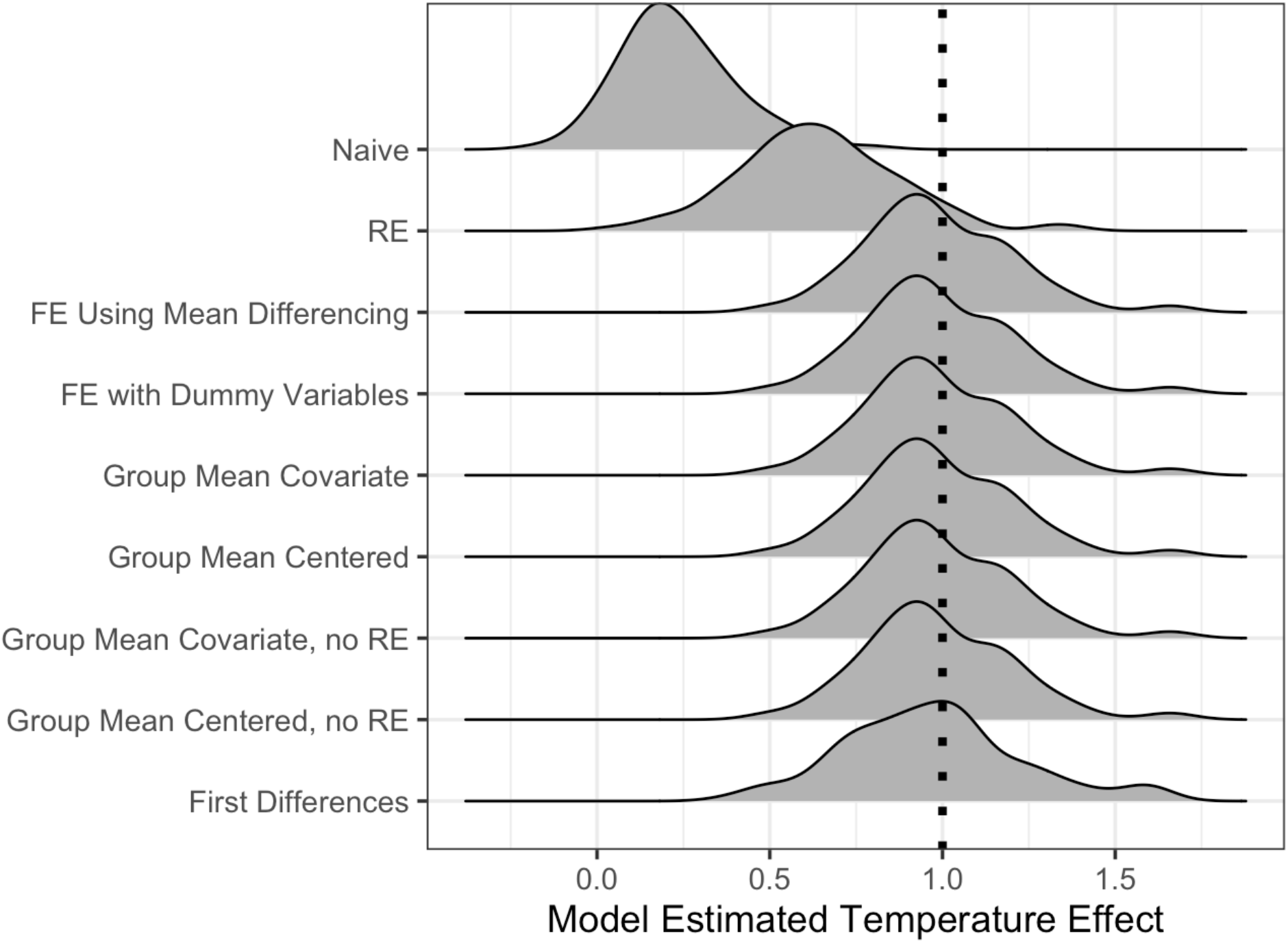
Distribution of point estimates of temperature effects from different models across all 100 simulations. The true effect size (= 1) is highlighted with a dotted line. The y-axis labels correspond to the Naïve model in equation 2, Random Effects (RE) model in equation 3, the Fixed Effects (FE) models in equations 5 and 6, the Group Mean Covariate models in equation 7 and the Group Mean Centered models to equation 8, and the First Differences model in the equation 9. The Naïve and Random Effects models produce biased coefficient estimates on average, in contrast to all other methods.

Unlike random effects in a mixed model design, *λ*_*i*_ is not constrained to be drawn from a predefined probability distribution. These designs will control for omitted variable bias from site-level observed and unobserved confounding variables and produce identical results to the preceding model for β_1_(Angrist & Pischke 2008; Wooldridge 2010), which we demonstrate with simulations below. While these two versions of the fixed effect design look different, they are equivalent (Angrist & Pischke 2008; Wooldridge 2010).

Fixed effect designs allow us to relax the strong assumption that all confounding variables are observed, measured, and included as covariates in models for a causal interpretation of β_1_when other assumptions are met (see Discussion). For ecological examples, see Larsen (2013), Dee et al. (2016), Dudney et al. (2021), Ratcliffe et al. (2023), and Dee et al. (2023).

Fixed effect designs have some drawbacks, despite their simplicity and strength in controlling for both observed and unobserved confounding variables. First, while these estimators make much weaker assumptions about confounding variables, they are inefficient compared to random effects. For each fixed effect (e.g., site), we estimate a separate coefficient. Estimating more parameters eats up degrees of freedom, requiring a larger sample size to achieve the same level of precision as random effects. This tradeoff creates a bias-variance trade-off (Bell *et al*. 2018). If one’s goal is causal inference, reducing bias is critical and thus fixed effects designs are preferable over a mixed-effects model (see simulation results in Table 1 and Fig. 6). Finally, with the fixed effects approach, we lose information about between-site variation, including gradients between sites that may be of interest. The fixed effects absorb this variation. These gradients, while confounded with other variables, could be the focus of some research questions which cannot be easily addressed using fixed effect designs.

**Table 1.** Summary of simulation results comparing estimates from each study design compared to a naïve bivariate correlation. Mean and SD of point estimates of temperature effects from different models in the first two columns. Fraction of simulated runs where the mean +/-2 SE of the temperature effect either overlapped 0 (i.e., high likelihood of committing a type II error) or did not contain the true effect of temperature in the final columns. Models are as in Fig. 7.

| Model Type | Mean Estimate | SD Estimate | Fraction Sims where 95% CI Contains 0 | Fraction Sims where 95% CI does Not Contain 1 |
| --- | --- | --- | --- | --- |
| Naïve | 0.231 | 0.165 | 0.56 | 0.99 |
| Random Effects (RE) | 0.640 | 0.232 | 0.08 | 0.54 |
| FE<sup>†</sup> Using Mean Differencing | 0.985 | 0.215 | 0.00 | 0.05 |
| FE<sup>†</sup> with Dummy Variables | 0.985 | 0.215 | 0.00 | 0.05 |
| Group Mean Covariate | 0.985 | 0.215 | 0.00 | 0.05 |
| Group Mean Centered | 0.985 | 0.215 | 0.00 | 0.05 |
| Group Mean Covariate, no RE | 0.985 | 0.215 | 0.01 | 0.04 |
| Group Mean Centered, no RE | 0.985 | 0.215 | 0.01 | 0.04 |
| First Differences | 0.971 | 0.259 | 0.01 | 0.12 |

<sup>†</sup>FE = econometric fixed effects

### Group Means for Efficiency, Inference, Fun, and Profit

What if we care about the between-site variation and comparisons across site are central to our question? To study between-site variation and mitigate the loss of efficiency from the fixed effect design, we can instead use **correlated random effects** designs (using terminology of Antonakis *et al*. 2021). Correlated random effects designs use group means of our causal variable of interest to control for confounding variables. For each cluster (e.g., each site, year), researchers calculate a group mean of the causal variable of interest (e.g. average temperature of a site) and include it as a group-level predictor. These group means of the causal variable control for the effects of confounders at the cluster level by acting as a proxy for confounders.

One correlated random effects design is the **Mundlak Device** (Mundlak 1978) and has many extensions (e.g., Wooldridge 2021). For clarity, we term it a **Group Mean Covariate** design, as in the following equation:

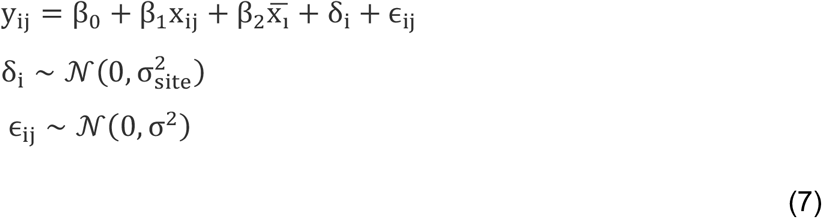

where 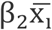, with 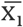, as the average site temperature accounts for the effect of cluster-level confounders and δ_i_ is a random effect of that cluster (i.e., site, other coefficients are as before or see Supplementary Table 1). From the path model in Figure 5C, we can see the site mean temperature is statistically controlled for in estimating the within-site temperature effect.

Using group means of our causal variable enables us to estimate a coefficient for between-cluster effects (e.g., between sites, β_2_), called a **contextual effect** (Antonakis *et al*. 2021). These coefficients contain a combination of causal and confounded effects and are not causally identified. They should not be taken as strong evidence supporting or refuting a particular hypothesis. The coefficient for the contextual effect, here site mean temperature, quantifies how changing the mean temperature of a site – and all properties that correlate with site mean temperature – affects snail abundance if temperature within a plot stayed the same. For example, *if our plot was 10 degrees C, what would snail abundance be if that plot was in a site with an average temperature of 5 degrees C versus 20 degrees C*? If the contextual effect is 0, we can conclude that a simple mixed-effects model would suffice and that omitted variable bias was not substantial in this particular analysis (Antonakis *et al*. 2021).

The Group Mean Covariate design will run into problems, however, if the correlation between our causal variable of interest and its cluster-level mean is too high. To overcome this issue, we can instead use a **Group Mean Centering** design, which transforms our causal variable to remove this correlation. Group Mean Centering subtracts the cluster-level mean from the causal variable of interest: 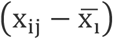. In our example, we subtract the site’s average temperature across the whole time series from the observed temperature for each year at each site (see Fig. 5D). After this transformation, we use this cluster-level centered variable (i.e., within cluster anomaly) as our predictor variable of interest with β_1_estimating the effect of a 1-degree change in anomaly. We control for cluster level mean – which includes confounding effects – as follows:

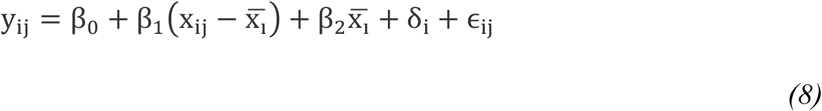

Equation 8 decomposes our causal variable of interest into between- and within-cluster terms, β_2_ and β_1_respectively. This is an approach already in use in ecology (van de Pol & Wright 2009). The coefficient of the site mean temperature, β_2_, is now the between-site effect of temperature and confounders. The coefficient, β_1_, is the within-site temperature effect and is the same as previous models (Supplementary Table 1). The interpretation of β_2_ is different than in the Group Mean Covariate design. β_2_ for our snail example is a **between estimator** of the combined effect of moving across gradients in temperature and correlated drivers between the sites but holding anomaly constant. For example, if we moved from a site with a 5ºC average temperature to one that was 10ºC, how would snail abundance change in a plot holding anomaly constant? If the anomaly was 1ºC, this would be a comparison between plots that were 6º at one site and 11ºC at another. But the result could be very different than comparing plots that were 6º and 11ºC from the same site. If β_2_ = β_1_, omitted variables are not meaningfully influencing snail abundances; both the between and within site differences are due solely to temperature or multiple confounders have cancelled one another out.

The Group Mean Covariate, Group Mean Centered, and Fixed Effects designs all differ in structure but will yield the same point estimates of β_1_(Fig. 6, Table 1) under most conditions and with balanced data (see simulations below and Wooldridge 2010). Mundlak (1978) showed that correlated random effects and fixed effects are algebraically identical in linear models and only differ in their inferences. Thus, one might ask: *which design should I use*? This decision depends on the structure and size of one’s data (e.g., how many coefficients do you have the power to estimate given your sample size) and the question of interest (e.g., are you interested in between-site differences?). For example, do you have many sites and are only interested in the causal effect of temperature? Fixed effects design. Do you want to know how plot-level snail abundance would change if the average site temperature changes, but plot temperature stays the same? Group Mean Covariate design. Do you want to understand the effects of temperature while examining the net effect of many variables shaping between-site gradients? Group Mean Centered design. Each design can further be extended to cases where the magnitude of the causal variable of interest’s effect is moderated by the level of confounding variables (i.e., an interaction between unobserved confounders and our causal variable of interest – see SI 3: A Difficult Slope: Omitted Variables that Cause Variation in the Magnitude of the Causal Effect).

Finally, for all of these designs, we note that accounting for serial correlation, heteroskedasticity, and clustering of the error through cluster robust standard errors or random effects at the cluster level are important for standard error estimation and thus inferences (Abadie *et al*. 2017; Cameron & Miller 2015). For more, see SI 4.

### What a Difference Differencing Makes

Our examples thus far have focused on unobserved confounding variables that vary across space (i.e., between sites) rather than time. Time can be difficult, as it can enter the picture in several different way. In the case of omitted confounders varying solely across time and not space, we can extend the frameworks presented above, using years rather than sites as clusters. If time-varying confounders are uniform across sites, then we can use an econometric fixed effect of time and an econometric fixed effect of space (a two-way fixed effect aka TWFE, Wooldridge 2021) and extensions (Roth *et al*. 2023) or a site-average of predictors and a time-average of predictors (a Two-Way Mundlak model design; Wooldridge 2021).

If, however, temporal confounders differ by site, we need a more general solution. If our causal variable of interest has the same trend over time as temporal confounders, we can use **temporal differencing**. For example, consider ocean warming alongside coastal development at our sites (Fig. S2) with development increasing over time, like warming, but varying in rate between sites. To separate these correlated drivers, we can use a **first-differences design** (Fig. 5e). We subtract all drivers and responses in year j from their value in year j-1 (i.e., Δ*y*_ji_ is change in snail abundance at site *i* between year *j* and *j-1*). We illustrate this showing the subtraction to produce the first differences model:

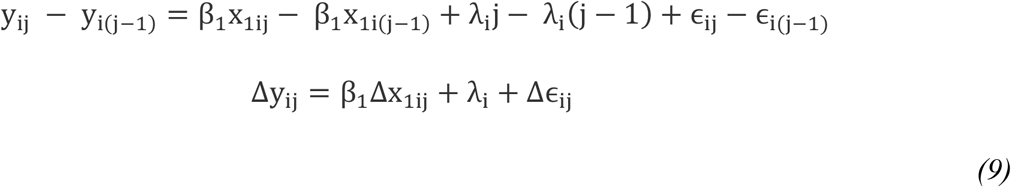

Here, *λ*_i_ is the slope of the site-specific temporal confounder. Spatial confounders are eliminated algebraically, just as in the fixed effects transformation. Finally, if we are uninterested in site-specific trends, we can calculate the **second difference** Δ^2^*y*_ji_=Δ*y*_ji_−Δ*y*_i,j-1_eliminating the need to estimate *λ*_i_ (Fig. 5f). While both designs control for temporal and spatial confounding, they cost one to two years of data. Finally, if omitted confounders vary spatiotemporally without trends or spatial and temporal confounders interact, we can extend the principles discussed here to more exotic designs (see Box 2).

### Comparison of Approaches

To demonstrate the utility the preceding solutions, and the consequences of not using them, we fit a variety of models to simulated data based on a longitudinal study of snail populations at multiple sites based on Figure 2. Snail abundance at site *i* in year *t* is a function of recruitment, temperature and other unobserved confounded drivers. For a single simulation run, we created a system as follows, simulating 10 sites over 10 years using a panel sampling design with:

- Oceanography as a random normal variable with a mean of 0 and standard deviation of 1. O_i_ ∼ N(0,1).
- Site mean recruitment calculated as -2 multiplied by the oceanography variable and then rescaled to have a mean of 10 individuals per plot (e.g., so it does not go negative). It is the same in a site across all years. R_i_ | O_i_ = -2 O_i_ + 10
- Site mean temperature as calculated as twice the oceanography variable and then rescaled to have a mean of 15C. T_i_ | O_i_ = 2 O_i_ + 15
- Site temperature in year *<u>j</u>* determined by site mean temperature and additional variation random normal variation with a mean of 0 and standard deviation of 1. T_ij_ ∼ N(T_i_, 1)
- Snail abundance at site *i* in is determined in a given year as in Fig. 2, as a function of recruitment, temperature and other drivers; the effects of both temperature and recruitment on snail abundance are 1. Other drivers are drawn from a random normal distribution with mean effect of 0 and standard deviation of 1. S_ij_ | T_ij_, R_i_ ∼ N(T_ij_ + Ri, 1)

Simulation code and results from 100 simulated data sets are in SI 6 and at https://github.com/jebyrnes/ovb_yeah_you_know_me. We analyzed each simulation run using all of the statistical designs above, including a naïve model with no site effect. We also included group mean covariate and group mean centered models with and without a random effect of site to demonstrate the role of a random effect in estimating standard errors and handling unbalanced data. Supporting information 7 uses Supporting Data S1 to analyze one data set showing the simplicity of implementation. For an interactive exploration of the full suite of simulated data and parameters, see the web applications in SI 8 (for one simulated run alone) and SI 9 (using replicated simulations to explore parameter distributions).

Our simulations show that the random effects (RE) model – what ecologists typically use – is consistently biased in these simulations. The point estimates from RE model are well-below the estimates from both the other designs and the true effect size (Fig. 7, Table 1). Not only is the estimated coefficient of the RE model always biased compared to other estimators in our simulations, it is more often within 2SE of 0 (i.e., would fail to reject a null hypothesis) in comparison to all other designs. More worrying, in the majority (54%) of simulations, the 95% confidence intervals of the RE model do not contain the true effect of 1 (Table 1). Other than the naïve and random effects model, the other designs show similar estimates with balanced data (in Table 1). The first differences model underperforms with respect to its CI not containing the true parameter value relative to other designs, but it performs far better than the RE and naïve design. However, the relative performance of first difference versus econometric fixed effect depends on the structure of data, whether it has more time periods, or more units (Wooldridge 2010), which likely explains this discrepancy.

Additional explorations show that, in line with the benefits of random effects in mixed models, a site-level random effect is crucial for Group Mean Centered or Group Mean Covariate models when either the study design is unbalanced or there is site-level variation that is uncorrelated with temperature (for more details, see SI 6). If our simulation has no site-level variation other than temperature and our confounder, a random effect does not improve either models’ ability to estimate the effect of our causal variable of interest with respect to bias or precision. This assumption is unrealistic for most real data sets, however. For estimating standard errors, in general, we urge researchers to incorporate random effects or clustered robust standard errors as needed to accommodate clustering and heteroskedasticity in the error, per the study design, recognizing the tradeoffs of using both and appropriate context (*reviewed in* Oshchepkov & Shirokanova 2022).

#### Box 2

**Reality Bites: Coping with Spatiotemporal Omitted Confounders**

Spatiotemporal confounding variables – those that are site (or plot) specific and vary through time – pose challenges, and the solutions require more thoughtful study designs. To illustrate, we consider a scenario where recruitment, a confounding variable related to both snail abundance and temperature, is not static through time but instead varies by site and year. For example, sites that experience strong cold-water pulses in a year also experience unusually snail high recruitment in those same years due to oceanographic drivers. The sampling designs for coping with spatiotemporal omitted variables are based on the same principles as cross-sectional and longitudinal sampling, only now we combine the two to include plots within sites that are sampled through time.

With longitudinal data with multiple plots sampled within a site through time, we can flexibly control for spatiotemporal confounding at the site level by extending the two-way fixed effect designs discussed above. We can add a site-by-time fixed effect, *η*_*ij*_, to our model, in addition to a fixed effect of plot, *λ*_k_, where *k* is a fixed plot within site resampled over time (see below for a discussion of fixed versus re-randomized plots). This produces the following means model (i.e., not using dummy coding but just the means for the fixed effects):

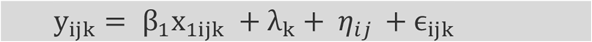

From this equation, we can see that *λ*_k_ captures time invariant plot-level confounding effects while *η*_*ij*_ captures the effects of spatiotemporal omitted variables at the site by time level. See Supplementary Table 1 for exact coefficient definitions. Note, there could be additional spatial or temporal only confounders. This design sweeps their effects onto the spatiotemporal term.

In small datasets, the above model design can consume degrees of freedom rapidly. In datasets with insufficient power, we can instead use the correlated random effects (e.g., a variation on the Two-way Mundlak model design *sensu* Wooldridge 2021) which are more efficient. Correlated random effect use site-year means 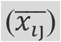 and plot means 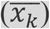 of temperature for the entire survey to control for spatiotemporal and plot confounding respectively:

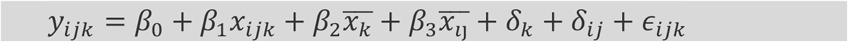

Here the *δ*_k_ and *δ*_*ij*_ terms are random effects for plot and unique site-time combinations respectively.

When sampling to handle spatiotemporal confounders, should plots within sites over time be permanent or randomly placed each year? The above models assume permanent plots, so we can eliminate confounding variables at the plot-level that is time invariant over the study period. For this reason, permanent plots help us cope with within-site OVB issues and have higher power to detect change over time (Urquhart & Kincaid 1999). Logistically, however, permanent plots within sites might not be possible. As such, the above models can be modified to drop plot effects; however, they would then assume that there are no confounding differences across plots and could have lower power to detect effects of drivers. We emphasize that the choice of fixed or random plot placement with these designs is a balancing act, however, as fixed plots can lead to a lower sample size due to logistical considerations in many environments, and direct readers to other explorations of this topic (e.g. Gomes 2022).

Finally, without variation within sites as well as through time – e.g., plots within sites resampled over years – we cannot include a site by year effect as in the above models. We can attempt to use site-level time trends (e.g., as linear or polynomial trends) as in Dee *et al*. 2016a or trends generated from Generalized Additive Models (Wood 2017) to approximate site-by-time, but this approach requires stronger assumptions about how confounders vary across sites over time. Therefore, we recommend researchers test the robustness of these assumptions. In cases without multiple plots per site over time, researchers can use sensitivity tests (Oster 2019) or partial identification (Ghassami *et al*. 2024; Miao *et al*. 2018) to get the bounds of estimates. In situations where these bounds are wide, however, without multiple plots per site, “nothing to be done” (Beckett 1954).

In general, we urge caution when dealing with spatiotemporal omitted variables, and careful use of causal diagrams to ensure that we are controlling for a confounder at the relevant spatiotemporal scale. This topic deserves far more exploration in Ecology.

## Discussion

We aim to introduce and spur broader uptake of techniques that address omitted variable bias for causal inference with observational data. At the core of these and other causal inference techniques is an *a priori* causal model of how a system works to guide sampling, statistical design choices, and to clarify assumptions required for a causal interpretation of estimated effects. Inferences made from designs that better control for unobserved confounding variables can improve our ability to understand biological systems, as seen in our simulations and results. Further, the techniques presented here are well within the standard statistical abilities of most modern ecologists (see SI 7 for R code).

We hope that Ecologists can see the concepts presented here as part of a generalizable approach to handling confounding variables using clustered or hierarchical data. While we use sites and years, the same concepts apply to studies with cohort effects, individual effects, or other lower levels of clustering as well as larger-scale studies with not just sites and years but regions and decades. Cross-sectional and longitudinal sampling designs are also generalizable beyond our example’s simple case, including for spatiotemporal designs (*see* Box 2). Combining these sampling designs with others, such as a stratified random sampling design (Foster *et al*. 2018; Grafström & Lundström 2013; Kermorvant *et al*. 2019; Robertson *et al*. 2013; Stevens & Olsen 2004), will allow for the analyses that can both improve causal identification and also provide more precision in estimation over multiple environmental gradients. How to design a study to fully account for confounders, however, will hinge on a causal structure of the system and a researcher’s ability to be humble in the face of what they might not know.

The approaches presented here are not a panacea. They too require assumptions for causal inference, as does any approach, including experiments (Kimmel *et al*. 2021). Some assumptions are shared with experiments: i.e., the Stable Unit Treatment Value Assumption (SUTVA) (reviewed in Kimmel *et al*. 2021). But these approaches require additional assumptions in terms of how they handle confounders (see Table S1). In addition, most of the statistical designs presented here include assumptions that effects are linear, additive (Imai & Kim 2021), and homogeneous (at least on the link scale in the case of Generalized Linear Models) across units and time periods. Generalized linear models may exhibit a slight downward bias for some methods, although this has been found to be largely negligible (Bell *et al*. 2018; Brumback *et al*. 2010) and techniques are being actively developed to ensure consistent estimation (Schunck & Perales 2017). We have included discussion of relaxing these assumptions via interactions (SI 3) and refer readers to a growing literature on estimating causal effects under more varied forms of heterogeneity and non-linearity (Callaway & Sant’Anna 2021; de Chaisemartin & D’Haultfœuille 2020; Goodman-Bacon 2021; Sun & Abraham 2021) or using flexible machine learning (Athey *et al*. 2019; Athey & Imbens 2019; Fink *et al*. 2023).

Further, the approaches presented here make the parallel trends assumption. The assumption of parallel trends can be understood considering a binary causal variable (i.e., if a driver is present or absent). It implies that, without driver being present, the *difference* in outcomes between different clusters (e.g., sites) after conditioning on covariates is constant through time. The assumption can be tested pre-treatment but is untestable after treatment (for details, see Roth 2022). The parallel trends assumption has come under a great deal of scrutiny (reviewed in Roth *et al*. 2023), particularly when changes in the causal variable of interest happen at different points in time across units (Baker *et al*. 2022; Marcus & Sant’Anna 2021) or with heterogeneous effects of causal variables (see Borusyak *et al*. 2023; de Chaisemartin & D’Haultfœuille 2020; Goodman-Bacon 2021; Sun & Abraham 2021). Solutions are evolving rapidly (*reviewed in* Roth et al. 2023), including for heterogeneous effects (*see* Roth *et al*. 2023), non-linear cases (Imai & Kim 2021), and continuous causal variables (Callaway *et al*. 2021) and are already being implemented in standard software (*reviewed in* Roth *et al*. 2023).

The important thing is to be transparent in how we deal – or do not deal – with assumptions required for causal interpretation of estimates, especially about confounding variables. What are the assumptions you are making to interpret an effect as causal? Why did you control for some covariates and not others? Do you have a DAG or even a conceptual model of your system that might help a reader understand your thought process? If you are using mixed models, do you meet the random effects assumption such that your clusters are not correlated with your predictor of interest, and why or why not? Have you evaluated your residuals to determine if you need to implement robust standard errors? Clarifying these and other decisions, even in a brief sentence if not a figure or full breakdown in a manuscript supplement (e.g., see Dee *et al*. 2023), will help make analyses more transparent and allow the work to be built on (e.g. by relaxing assumptions) to advance science. Further, in concert with the approaches presented here, we suggest using sensitivity tests (Altonji *et al*. 2005; Cinelli & Hazlett 2020; Oster 2019; Rosenbaum 2002) and other designs that differ in assumptions to assess robustness of results (*see* Dee *et al*. 2023 for an ecological example). At the end of the day, we must be humble and accept that our models and knowledge are imperfect. Someday, someone will come along with a different approach or new data that produces different conclusions and yields new insights, which is just part of the scientific process.

Finally, we emphasize that this paper provides an entry point into a broader, interdisciplinary literature on causal inference (see SI 5 for useful texts). Other quasi-experimental designs, such as instrumental variables, synthetic control, and regression discontinuity, can also be used to eliminate omitted variable bias (see Arif & MacNeil 2022b; Butsic *et al*. 2017; Dee *et al*. 2023; Fick *et al*. 2021; Grace 2021; Kendall 2015; Larsen *et al*. 2019; MacDonald & Mordecai 2019). Thoughtful uses of the front-door criterion – using mediators between a cause and effect unaffected by confounders – could also prove useful for causal analysis (Bellemare *et al*. 2024; Pearl *et al*. 2016); although, there are none to our knowledge in the Ecological literature yet. We urge ecologists, long grounded in experiments, to consider these and other transdisciplinary advances in causal inference in observational data – as an important complement to experiments.

## Conclusion

“Correlation does not equal causation” rings in many of our heads from Biostatistics 101. A key reason behind this message is the specter of Omitted Variable Bias. This fear has impeded the use of observational data for causal inference in Ecology for much of its recent history. We hope this review can lift some of that fear and, armed with the new tools and knowledge of a literature beyond this piece (see above), we can move forward as a discipline. With a massively growing volume of observational data, and problems at continental to global scales demanding rapid answers, we look forward to seeing Ecologists harness these techniques to answering crucial questions on into the future.

## Acknowledgements

We thank the NCEAS Long-Term Ecological Research (LTER) working group, “Scaling-up productivity responses to changes in biodiversity,” for initiating the conversations that led to this paper, supported by the National Science Foundation as (NSF) LTER Network Communications Office and DEB-1545288. This work was partially supported by the NSF as part of the PIE-LTER Program (award #1637630), Woods Hole Sea Grant, and the Stone Living Lab to J.B.; and by NSF OCE #2049360, NSF CAREER #2340606, and NASA BioSCape award #80NSSC22K0796 to L.E.D. We thank three anonymous reviewers for comments on the submitted manuscript and comments from readers of our biorXiv preprint.We thank S. Elmendorf, B. Hobart, C. Severen, I. Rosenthal, R. Stevenson, A. Carter, S. Miller, the UMB Causal Inference reading group, and the UMB Stats Snack for helpful conversation on early drafts.

