## Supplemental Tables, Figures, Text, Data, and Code for "Causal inference with observational data and unobserved confounding variables": byrnes_dee_revision_supporting_s1-s5_and_tables.pdf

### <u>Contents</u>

**Supporting Table S1:** Models, equations, and the definition of coefficients based on linear regression models.

**Supporting Table S2:** Key assumptions, drawbacks, and types of Omitted Variable Bias handled by statistical model designs in this paper.

**Supporting Information S1.** Additional details and examples of Directed Acyclic Graphs and confounding

**Supporting Figure S1.** Examples of statistical control for confounding variables informed by causal graphs.

**Supporting Information S2.** Directed Acyclic Graphs: What about feedbacks?

**Supporting Information S3.** A Difficult Slope: Omitted Variables that cause variation in the magnitude of the causal effect

**Supporting Information S4.** Clustered Robust Standard Errors: An Underutilized Tool in Ecology

**Supporting Figure S2.** Directed Acyclic Graph of a scenario with spatial and temporal omitted variables where site-level temporal confounders are correlated with the causal variable of interest.

**Supporting Information S5**. Additional recommendations for readings

**Supporting Information S6.** Models and Simulations to Evaluate the Consequences of Model Structure for Omitted Variable Bias

**Supporting Information S7.** Implementing OVB Model Methods in R

**Supporting Data S1:** Data for analysis in supporting information S7.

**Supporting Information S8.** Code for R Shiny App to simulate a data set and compare results of statistical designs over just that data set.

**Supporting Information S9.** Code for R Shiny App to simulate many data sets and compare aggregate results of statistical designs.

**Supporting Table S1: Models, equations, and the definition of coefficients based on linear regression models.** For all models, *i* refers to site and *j* refers to year or plot, for longitudinal or cross-sectional designs, respectively, save spatiotemporal models where *j* is time and *k* is plot. $y_{ij}$ is the number of snails at site *i* in year or plot *j*. $x_{ij}$ is temperature at site *I* in year or plot *i*. $\bar{y_i}$ is average number of snails at site *i* over a whole study (i.e., either across the entire time period for a panel design or across an entire site for a cross-sectional design), and $\bar{x_i}$ is average temperature at site *i* similarly over a whole study.

Several designs include random effects, in which random effects are assumed to follow a normal distribution. We note random effects are estimated as part of the error, not as a parameter directly estimated (in comparison to the econometric fixed effect models). These model structures can also be modified for use in Generalized Linear Models and Generalized Linear Mixed Effects Models but are presented for the Linear Models and Linear Mixed Models for simplicity.

| Model | Equation(s) | Terms |
|---|---|---|
| Naïve Model | $y_{ij} = \beta_0 + \beta_1 x_{ij} + \epsilon_{ij}$<br>$\epsilon_{ij} \sim \mathcal{N}(0, \sigma^2)$ | $\beta_0$: Average number of snails when temperature is 0.<br>$\beta_1$: Increase in snails per 1 degree increase in temperature.<br>$\epsilon_{ij}$: Other drivers of snail abundance, random noise, and measurement error. |
| Random Effects Model | $y_{ij} = \beta_0 + \beta_1 x_{ij} + (\delta_i + \epsilon_{ij})$<br>$\delta_i \sim \mathcal{N}(0, \sigma_{site}^2)$<br>$\epsilon_{ij} \sim \mathcal{N}(0, \sigma^2)$ | $\beta_0$: Average number of snails when temperature is 0 across all sites.<br>$\beta_1$: Increase in snails per 1 degree increase in temperature.<br>$\delta_i$: The random effect. Other drivers of snail abundance, random error, and measurement error at the site level across the study.<br>$\epsilon_{ij}$: Other drivers of snail abundance, random noise, and measurement error. |
| Fixed Effects Transformation (Within Transformation with de-meaning) | $y_{ij} - \bar{y_i} = \beta_1(x_{ij} - \bar{x_i})$<br>$+(\epsilon_{ij} - \bar{\epsilon_i})$ | $\beta_1$: Increase in snails per 1 degree increase in temperature, holding all other covariates constant. Note:all site-level covariates, including temperature, are held at at zero after demeaning, or their mean value before demeaning.<br>$\bar{\epsilon_i}$: Site-level average of other drivers of snails, random error, and measurement error. |

| Model | Equation(s) | Terms |
|---|---|---|
| Fixed Effects Means Model | $y_{ij} = \beta_1 x_{ij} + \lambda_i + \epsilon_{ij}$<br>$\epsilon_{ij} \sim \mathcal{N}(0, \sigma^2)$ | $\beta_1$: Increase in snails per 1 degree increase in temperature, holding all other covariates constant.<br>$\lambda_i$: Site-level, time or plot invariant effects. The average number of snails at site *i* when all site-level covariates, including temperature, are held at zero.<br>$\epsilon_{ij}$: Other drivers of snail abundance, random noise, and measurement error. |
| Group Mean Covariate Model (aka Mundlak Device) | $y_{ij} = \beta_0 + \beta_1 x_{ij} +$<br>$\beta_2 \bar{x_i} + \delta_i + \epsilon_{ij}$<br>$\delta_i \sim \mathcal{N}(0, \sigma^2_{site})$<br>$\epsilon_{ij} \sim \mathcal{N}(0, \sigma^2)$ | $\beta_0$: Average number of snails when temperature is 0 and average site temperature is 0.<br>$\beta_1$: Increase in snails per 1 degree increase in temperature, holding all other covariates constant.<br>$\beta_2$: Change in snail abundance given a 1 degree increase in average site temperature and all other site-level confounding variables, holding temperature constant.<br>$\delta_i$: The random effect of site. Other drivers of snail abundance, random error, and measurement error at the site level across the study.<br>$\epsilon_{ij}$: Other drivers of snail abundance, random noise, and measurement error. |
| Group Mean Centered Model | $y_{ij} = \beta_0 + \beta_1 (x_{ij} - \bar{x_i}) +$<br>$\beta_2 \bar{x_i} + \delta_i + \epsilon_{ij}$<br>$\delta_i \sim \mathcal{N}(0, \sigma^2_{site})$<br>$\epsilon_{ij} \sim \mathcal{N}(0, \sigma^2)$ | $\beta_0$: Average number of snails when temperature is 0 and the average site temperature is 0.<br>$\beta_1$: Increase in snails per 1 degree increase in temperature, holding all other covariates constant.<br>$\beta_2$: Change in abundance of snails moving to a site that is 1 degree warmer on average, along with confounded drivers, holding temperature anomaly constant.<br>$\delta_i$: The random effect of site. Other drivers of snail abundance, random error, |

| Model | Equation(s) | Terms |
|---|---|---|
| | | and measurement error at the site level across the study.<br>$\epsilon_{ij}$: Other drivers of snail abundance, random noise, and measurement error. |
| First Differences Model | $\Delta y_{ij} = \beta_1 \Delta x_{ij} + \lambda_i + \Delta\epsilon_{ij}$<br>$\Delta\epsilon_{ij} \sim \mathcal{N}(0, \sigma^2)$ | $\Delta y_{ij}$ and $\Delta x_{ij}$: Change in snails and temperature at one site between two time points.<br>$\beta_1$: Increase in snails per 1 degree increase in temperature, holding all other covariates constant. Note: all site-level covariates that do not change over time, including temperature, are held at zero by calculation of first differences.<br>$\lambda_i$: Site-level slope of any confounded drivers that are changing over time alongside temperature.<br>$\Delta\epsilon_{ij}$: Change in other drivers of snail abundance, random noise, and measurement error. |
| Second Differences Model | $\Delta y_{ij}^2 = \beta_1 \Delta^2 x_{ij} + \Delta^2 \epsilon_{ij}$<br>$\Delta\epsilon_{ij}^2 \sim \mathcal{N}(0, \sigma^2)$ | $\Delta^2 y_{ij}$ and $\Delta^2 x_{ij}$: Change in the change in snails and temperature at one site between two time points. I.e., $\Delta y_{i1} - \Delta y_{i2}$.<br>$\beta_1$: Increase in rate of change in snails per 1 unit increase in rate of change in temperature, holding all other covariates constant. Note: all site-level covariates, including temperature, are held at zero after calculating second differences save those that are change in a non-linear fashion over time.<br>$\Delta^2 \epsilon_{ij}$: Acceleration in other drivers of snail abundance, random noise, and measurement error. |
| Two-Way Fixed Effects Means Model (TWFE) | $y_{ij} = \beta_1 x_{ij} + \lambda_i + \eta_j + \epsilon_{ij}$<br>$\epsilon_{ij} \sim \mathcal{N}(0, \sigma^2)$ | $\beta_1$: Increase in snails per 1 degree increase in temperature, holding all other covariates constant.<br>$\lambda_i$: Site-level, time or plot invariant effects. The average number of snails at |

| Model | Equation(s) | Terms |
|---|---|---|
| | | site $i$ if temperature was 0 without considering year effects.<br>$\eta_j$: Year-level site invariant effects. The average number of snails in year $j$ if temperature was 0 without considering site effects.<br>$\epsilon_{ij}$: Other drivers of snail abundance, random noise, and measurement error. |
| Two-Way Group Mean Covariate Model (Mundlak Device) | $y_{ij} = \beta_0 + \beta_1 x_{ij} + \beta_2 \bar{x_i} + \beta_3 \bar{x_j} + \delta_i + \epsilon_{ij}$<br>$\delta_i \sim \mathcal{N}(0, \sigma^2_{site})$<br>$\epsilon_{ij} \sim \mathcal{N}(0, \sigma^2)$ | $\beta_0$: Average number of snails when temperature is 0 and average site temperature is 0.<br>$\beta_1$: Increase in snails per 1 degree increase in temperature, holding all other covariates constant.<br>$\beta_2$: Change in snail abundance given a 1 degree increase in average site temperature and all other site confounded drivers, holding temperature constant.<br>$\beta_3$: Change in snail abundance given a 1 degree increase in average temperature and changes in associated temporal confounders in a year, holding temperature constant.<br>$\delta_i$: The random effect of site. Other drivers of snail abundance, random error, and measurement error at the site level across the study.<br>$\epsilon_{ij}$: Other drivers of snail abundance, random noise, and measurement error. |
| Heterogeneous Group Mean Covariate Model | $y_{ij} = \beta_0 + \beta_1 x_{ij} + \beta_2 \bar{x_i} + \beta_3 x_{ij} \bar{x_i} + \delta_i + \epsilon_{ij}$<br>$\delta_i \sim \mathcal{N}(0, \sigma^2_{site})$<br>$\epsilon_{ij} \sim \mathcal{N}(0, \sigma^2)$ | $\beta_0$: Average number of snails when temperature is 0 and average site temperature is 0.<br>$\beta_1$: Increase in snails per 1 degree increase in temperature.<br>$\beta_2$: Change in snail abundance given a 1 degree increase in average site temperature and all other site confounded drivers, holding temperature constant. |

| Model | Equation(s) | Terms |
|---|---|---|
| | | $\beta_3$: Change in effect of temperature per 1 degree change in site mean average temperature.<br>$\delta_i$: The random effect of site. Other drivers of snail abundance, random error, and measurement error at the site level across the study.<br>$\epsilon_{ij}$: Other drivers of snail abundance, random noise, and measurement error. |
| Spatio-Temporal Fixed Effects Model | $y_{ijk} = \beta_1 x_{1ijk} + \alpha_k + \eta_{ij} + \epsilon_{ijk}$<br>$\epsilon_{ijk} \sim \mathcal{N}(0, \sigma^2)$ | $\beta_1$: Increase in snails per 1 degree increase in temperature.<br>$\alpha_k$: Plot-level fixed effect for plot *k* in site *i*. The average number of snails in plot *k* when all plot-level covariates, including temperature, are held constant without considering site-time effects.<br>$\eta_{ij}$: Site-time fixed effects. The average number of snails at site *i* in year *j* when all site-by-year effects (including site and year specific temperature) are held constant without considering plot effects.<br>$\epsilon_{ijk}$: Other drivers of snail abundance, random noise, and measurement error. |
| Spatio-Temporal Group Mean Covariate Model | $y_{ijk} = \beta_0 + \beta_1 x_{ijk} + \beta_2 \bar{x_k} + \beta_3 \bar{x_{ij}} + \delta_k + \delta_{ij} + \epsilon_{ijk}$<br>$\delta_k \sim \mathcal{N}\left(0, \sigma^2_{plot}\right)$<br>$\delta_{ij} \sim \mathcal{N}(0, \sigma^2_{site-time})$<br>$\epsilon_{ijk} \sim \mathcal{N}(0, \sigma^2)$ | $\beta_0$: Average number of snails when temperature is 0 and average site temperature is 0.<br>$\beta_1$: Increase in snails per 1 degree increase in temperature.<br>$\beta_2$: Change in snail abundance given a 1 degree increase in average plot temperature and all other plot confounded drivers, holding temperature constant.<br>$\beta_3$: Change in snail abundance given a 1 degree increase in average temperature and changes in associated site-time-level confounders, holding temperature constant.<br>$\delta_k$: The random effect of plot. Other drivers of snail abundance, random error, |

| Model | Equation(s) | Terms |
| --- | --- | --- |
| | | and measurement error at the plot within site level across the study.<br>$\delta_{ij}$: The random effect of site-time. Other drivers of snail abundance, random error, and measurement error at a site within a year.<br><br>$\epsilon_{ijk}$: Other drivers of snail abundance, random noise, and measurement error. |

**Supporting Table S2. Key assumptions, drawbacks, and types of Omitted Variable Bias handled by statistical model designs in this paper**. We highlight causal assumptions related to confounding, rather than statistical assumptions (e.g., about functional form specification, error structure). All have the designs make the Stable Unit Treatment Assumption, which includes: 1) no interference and 2) no hidden and multiple versions of the treatment. We also do not discuss other sources of bias herein (i.e., from measurement error) or different possible forms of error and standard error estimation.

| **Model** | **Forms of OVB dealt with** | **Key Causal Assumptions** | **Drawbacks** |
|---|---|---|---|
| Naïve | None | • No confounding variables.<br><br>• Measurements within sites are independent.<br><br>• Homogeneous effects of causal variable of interest. | Biased estimator of $\beta_1$ in the presence of any confounding variables.<br><br>Measurements within site are likely not independent, regardless of confounding. |
| Random Effects | None | • No confounders that are unobserved and not included in the model.<br><br>• Homogeneous effects of causal variable of interest. | While it appears to incorporate a site effect, if there are other confounding drivers at the site level, we violate the assumption of endogeneity and create omitted variable bias. Thus, the estimate of $\beta_1$ is biased and not causal. |
| Fixed Effects Transformation | Site-level OVB removed via removing the mean value of site for both the predictor and response | • Confounders only at the site level, reflecting confounding differences between sites.<br><br>• Homogeneous effects of causal variable of interest.<br><br>• The difference between sites, after controlling for the causal driver and other | Removes between site variation. Not able to calculate counterfactual predictions. |

| Model | Forms of OVB dealt with | Key Causal Assumptions | Drawbacks |
|---|---|---|---|
| | | covariates, is constant (Parallel Trends assumption) | |
| Fixed Effects Models | Site-level OVB controlled for by site-level fixed effect. | • Confounders only operating between sites.<br><br>• Homogeneous effects of causal variable of interest.<br><br>• The difference between sites, after controlling for the causal driver and other covariates, is constant (Parallel Trends assumption) | Does not model between site variation.<br><br>Less efficient with many sites.<br><br>Not able to calculate counterfactual predictions about new sites. |
| Group Mean Covariate Model (aka Mundlak Device) | Site-level OVB controlled for by group mean term in model. This term encompasses the effect of both temperature at the site-level as well as other correlated site-level confounders. | • Confounders only operating between sites.<br><br>• Homogeneous effects of causal variable of interest.<br><br>• The difference between sites, after controlling for the causal driver and other covariates, is constant (Parallel Trends assumption) | Can get complex if multiple drivers are included in the model (i.e., need a group-mean for each).<br><br>Between site terms should be interpreted with caution, as they represent the effects of the driver and its confounders combined. |
| Group Mean Covariate Centered Model | Site-level OVB controlled for by group mean term in model. This term encompasses the effect of both temperature at the site-level as well as other correlated site-level confounders. | • Confounders only operating between sites.<br><br>• Homogeneous effects of causal variable of interest.<br><br>• The difference between sites, after controlling for the causal driver and other covariates, is constant (Parallel Trends assumption) | Can get complex if multiple drivers are included in the model (i.e., need a group-mean for each).<br><br>Between site terms should be interpreted with caution, as they represent the effects of the driver and its confounders combined. |
| First Differences Model | Site-level OVB removed via | • Confounders only operating between sites or | Evaluates effect of driver on change, but |

| Model | Forms of OVB dealt with | Key Causal Assumptions | Drawbacks |
|---|---|---|---|
| | examining change in driver and response rather than measured values alone. Site-level temporal confounders accounted for by additional site-level coefficient representing slope of site-level confounder exhibiting same temporal trend as causal variable of interest. | with same temporal trend of driver of interest within a site.<br><br>• Effect of temporal confounders is linear.<br><br>• Homogeneous effects of causal variable of interest.<br><br>• The difference between sites, after controlling for the causal driver and other covariates, is constant (Parallel Trends assumption). | not on observed value. Thus, cannot model the effect of a driver on the standing stock of a response variable at a site.<br><br>Loses one year of data for each site. Potentially inefficient if many sites by few years per site. |
| Second Differences Model | Site-level OVB and effects of site-level confounders with same temporal trend as driver of interest removed via examining acceleration in driver and response rather than measured values or change in both. | • Confounders only operating between sites or with same temporal trend of driver of interest within a site.<br><br>• Homogeneous effects of causal variable of interest.<br><br>• The difference between sites, after controlling for the causal driver and other covariates, is constant (Parallel Trends assumption). | Evaluates effect of driver on acceleration of change, but not on observed value. Thus, cannot model the effect of a driver on the standing stock of a response variable at a site.<br><br>Loses two years of data for each site.<br><br>Acceleration can be difficult to interpret as opposed to rate of change. |
| Two-Way Fixed Effects Model (TWFE) | Site-level and Year-level OVB controlled for by site- and year-level fixed effects. | • Confounders only operating between sites or between years. No spatio-temporal confounding.<br><br>• Homogeneous effects of causal variable of interest.<br><br>• The difference between sites, after controlling for | Does not model between site and between year variation.<br><br>Less efficient. With many years and sites, requires a high sample size. |

| Model | Forms of OVB dealt with | Key Causal Assumptions | Drawbacks |
|---|---|---|---|
| | | the causal driver and other covariates, is constant (Parallel Trends assumption). | Not able to calculate counterfactual predictions about new sites or years. |
| Two-Way Group Mean Covariate Model (Mundlak Device) | Site- and year-level OVB controlled for by group mean terms in model. | • Confounders only operating between sites or between years.<br><br>• Homogeneous effects of causal variable of interest.<br><br>• The difference between sites, after controlling for the causal driver and other covariates, is constant (Parallel Trends assumption). | Model complexity and higher sample sizes/range of variation in predictors needed to separate the effects of both spatial and temporal confounders via a group mean term.<br><br>Between site terms should be interpreted with caution, as they represent the effects of the driver and its confounders combined. |
| Heterogeneous Group Mean Covariate Model | Site-level OVB controlled for as well as how site-level confounders can alter a causal effect. | • Confounders only operating between sites.<br><br>• The difference between sites, after controlling for the causal driver and other covariates, is constant (Parallel Trends assumption). | Can get complex if multiple drivers are included in the model (i.e., need a group-mean for each). For high precision, requires high within-site variation in causal variable of interest to create an interaction term with sufficient range to differentiate from component parts.<br><br>Between site terms should be interpreted with caution, as they represent the effects of the driver and its confounders combined. |

| Model | Forms of OVB dealt with | Key Causal Assumptions | Drawbacks |
|---|---|---|---|
| Spatio-Temporal Fixed Effects Model | Plot-level and site by time level OVB controlled for by fixed effects terms in the model. | • Homogeneous effects of causal variable of interest. | Model requires multiple fixed plots per site sampled over time at multiple sites.<br><br>Number of parameters is very large, so the estimator is inefficient. |
| Spatio-Temporal Group Mean Centered Model | Plot-level and site by time level OVB controlled for by group mean terms in the model. | • Homogeneous effects of causal variable of interest. | Model requires multiple fixed plots per site sampled over time at multiple sites.<br><br>Between site terms should be interpreted with caution, as they represent the effects of the driver and its confounders combined. |

**Supporting Information S1. Additional details and examples of Directed Acyclic Graphs and confounding**

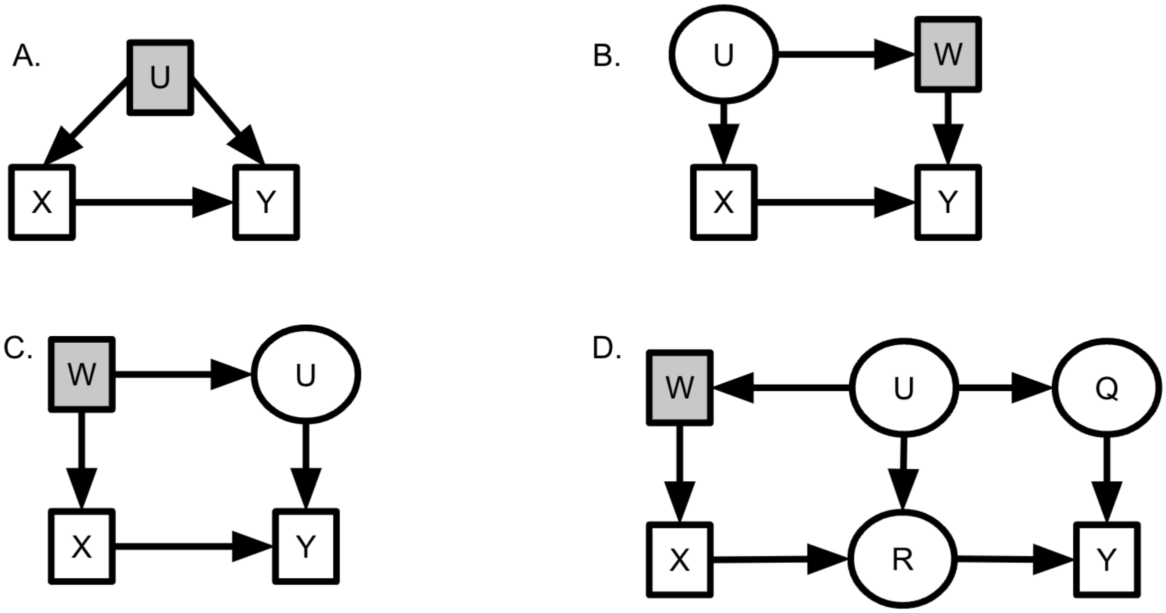


**Supporting Figure S1. Examples of statistical control for confounding variables informed by causal graphs.** By including shaded observed variables, either *U* or *W*, in a statistical analysis of the effects of *X* on *Y*, omitted variable bias is controlled for the results have a causal interpretation. The relationship between the control variable and *Y* might (as in panel **A** and **B**) or might (as in panel **C** and **D**) not have a causal interpretation, depending on the structure of the system. Note, in (**D**), *Q* would have also served as an adequate control instead of *W*. *R* would have been a bad control.

In the absence of a randomized experiment, one way to eliminate confounding from a measured variable (i.e., *U* in Fig. S1) is to include a variable in a statistical model. Including the variable controls for its confounding effect, thereby blocking all paths between *X* and *Y* via *U*. This variable to control for could be the confounder itself or another variable that is a "child node" or downstream of U that blocks the path affecting *X* or *Y* (e.g., *W* in Fig. S1 – if there was more than one path and *W* didn't block both, we'd need to control for a variable on the second path as well). Controlling for such a confounding variable means that the ensuing analysis will satisfy the **back-door criterion** (Pearl 1995, Fig. S1A); when all back doors are "shut", the influence of confounders are removed (i.e. we have established conditional independence between the causal variable of interest and the error term). Without including and thus controlling for these confounding variables or others that block their influence in a regression analysis (e.g. see Fig S1), omitted confounders will cause omitted variable bias.

**Supporting Information S2. Directed Acyclic Graphs: What about feedbacks?**

A common critique is that DAGs do not include feedbacks, to which we respond by asking the reader to think of their definition of causality. Here, we adopt the Neyman-Rubin counterfactual causality framework (Holland 1986; Rubin 1974, 2005) where we recognize that cause temporarily precedes effect. Therefore, feedbacks can be handled by thinking about a system with a temporal lag (e.g., Larson *et al.* 2008). If an instantaneous feedback is truly present, or if a time-series of both the driver and response variable is not available, one will likely require other tools such as instrumental variables - something beyond the scope of this manuscript (for a comprehensive review see Imbens 2014; for an ecological perspective see Grace 2021; for examples, MacDonald & Mordecai 2019; Dee *et al.* 2023).

**Supporting Information S3: A difficult slope: Omitted variables that cause variation in the magnitude of the causal effect**

An omitted confounder might not merely contaminate our estimate of a causal effect but can also lead to model misspecification in the form of missed heterogeneity in the causal effect. This occurs when the causal effect of our variable of interest depends on the level of the confounder itself (i.e., it modifies the causal effect – an interaction effect). In our example, thermal effects on snail abundance could depend on levels of recruitment because dense aggregations of intertidal organisms are often better at retaining water and thus resisting desiccation or other forms of thermal stress (e.g., Silliman *et al.* 2011). This dependence is problematic if we have not measured recruitment. In a naive mixed model, we might incorporate this heterogeneity as a random slope. As before, however, the random effects assumption is violated, so a random effects estimator will be biased. To deal with the problem of omitted variable bias in this case, we present two solutions. First, we can use a fixed effects design and include an interaction term between the site dummy variable and our causal variable of interest, allowing us to estimate site-specific temperature effects. Given that we now have site-level slopes, the number of parameters can blow up, leading to this approach being highly inefficient and not advisable for small sample sizes. Instead, we could use correlated random effects approaches with an interaction between the group mean and our causal variable of interest. For example, for a group mean covariate design (i.e. Mundlak device), we would use the following equation:

$$y_{ij} = \beta_0 + \beta_1 x_{ij} + \beta_2 \overline{x_i} + \beta_3 x_{ij}\overline{x_i} + \delta_i + \epsilon_{ij}$$

This design allows us to examine how site-level confounders – known and unknown – can lead to variation in the effect of our causal variable of interest. It can also show that they have no effect if the estimand for $\beta_3$ is not different from 0. We could use a similar model for the group mean centered design if deemed appropriate. If we suspect that the magnitude of the temperature effect varied with other non-confounded covariates, we could instead use a random slope. In general, models with interactions representing moderators can provide powerful insights into both the effect of the causal driver of interest as well as how those effects vary.

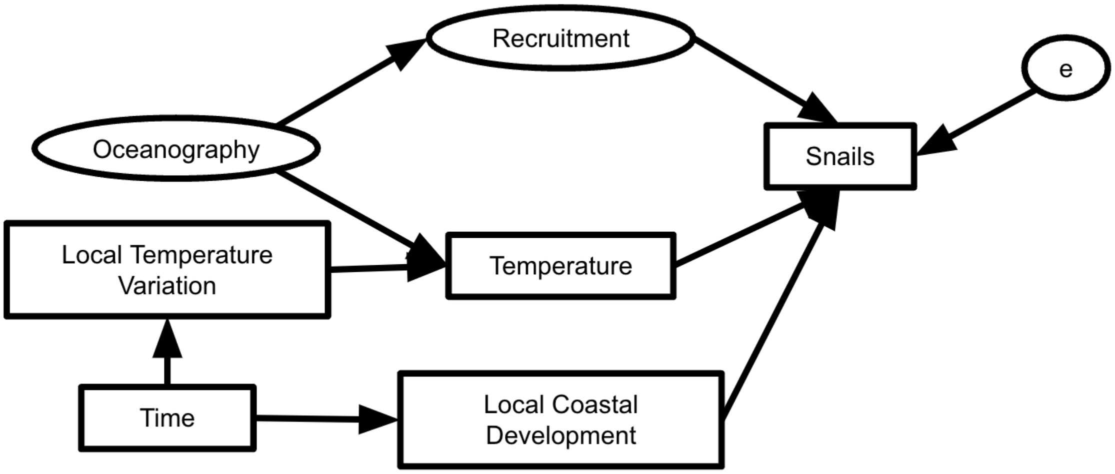


**Supporting Figure S2. Directed Acyclic Graph of a scenario with spatial and temporal omitted variables where site-level temporal confounders are correlated with the causal variable of interest.** The causal diagram shows the true data generating process of this hypothetical system, with not only recruitment confounded with temperature, but also with local coastal development increasing over time alongside local temperature also increasing over time. Thus, we have both temporal and spatial confounding variables which can be controlled for by first and second difference designs.

**Supporting Information S4: Clustered robust standard errors: An underutilized tool in Ecology**

While the focus of this paper is on bias not precision, many of the issues discussed overlap with issues of non-independence that could generate incorrect standard errors and statistical tests for inference. In light of that, we recommend the use of clustered robust standard errors. Clustered robust standard errors offer a flexibly way to accommodate clustered data, heteroskedasticity, serial correlation between time points, and other arbitrary correlation structures within the data (Abadie *et al.* 2017; Cameron & Miller 2015) yet are not commonly used in Ecology (but see examples in Dee *et al.* 2016; Dudney *et al.* 2021; and code in the appendices of Dee *et al.* 2023). While random effects, autocorrelation structures, and more, can address some of the same issues, clustered robust standard errors make weaker assumptions about the exact form of the correlation structures in the error, thus providing a simpler solution when the structure of the data are not of interest to the research question but important to account for in inferences. However, there are tradeoffs – weaker assumptions also mean less efficiency. Thus, we recommend looking at comparisons of approaches (e.g., Oshchepkov & Shirokanova 2022) or conducting sensitivity tests to check the robustness of inferences to choices of approach for modeling standard errors. A full discussion of clustered and robust standard errors is beyond the scope of this paper, and we refer applied researchers to the documentation for the '*sandwich'* package in R and other comprehensive reviews (e.g., Cameron & Miller 2015).

**Supporting Information S5. Additional recommendations for readings**

The field of causal inference spans disciplines, including Econometrics, Epidemiology, Computer Science, Public Health, and more that are developing rigorous approaches for causal inference in observational and experimental data. Embracing this transdisciplinary approach will enable us to enhance our knowledge of the tremendous advances in causal inference and explore questions currently beyond our reach. As an incomplete set of starting points for further reading, we recommend The Effect: An Introduction to Research Design and Causality (2021), Cunningham's Causal Inference: The Mixtape (2021), McElreath's chapters on causal diagrams in Statistical Rethinking (2020), Angrist and Pishke's Mostly Harmless Econometrics (2008),

Morgan and Winship's Counterfactuals and Causal Inference (2015), Sloman's Causal Models (2005), and Pearl's Causal Inference in Statistics: A Primer (2016).
