## Supplemental Tables, Figures, Text, Data, and Code for "Causal inference with observational data and unobserved confounding variables": supporting_information_s6.pdf

### Supporting Information S6: Models and Simulations to Evaluate the Consequences of Model Structure for Omitted Variable Bias

#### The System and the Study

To evaluate the impact of Omitted Variable Bias (OVB) on different models, consider a system where oceanography drives site temperature and recruitment over time. Temperature also fluctuates over time within each site. Recruitment and temperature influence snail abundance, as do other uncorrelated drivers. You have conducted a study in this system measuring 10 sites, each site sampled once per year over 10 years. In this study, you have recorded snail abundance and temperature. But have no measure of recruitment. Note, the results of models will be the same if you had instead sampled in one year across 10 sites with 10 plots per site if there were plot-level drivers of temperature that behaved the same as below.

To parameterize our simulations, consider the following:

- Our Oceanography latent variable has a mean of 0 and a SD of 1.
- Site temperature is twice the oceanography variable and transformed to have a mean of 15C.
- Site recruitment is -2 multiplied by the oceanography variable and transformed to have a mean of 15 individuals per plot.
- There is additional random variation between sites with a mean of 0 and SD of 1.
- Sites are sampled over 10 years.
- Within a site, the temperature varies over time according to a normal distribution with a mean of 1.
- There is a 1:1 relationship between temperature and snail abundance and recruitment and snails.
- Other non-correlated drivers in the system influence snail abundance with a mean influence of 0 and a SD of 1.

Thus, the system looks like this:

#### Functions to Create The System

To simulated data, let's begin by loading some libraries

```
library(tidyverse)
library(lme4)
library(broom)
library(broom.mixed)
library(DiagrammeR)
library(glue)

theme_set(theme_bw(base_size = 14))
```

Next, we need a function that will create a template of simulated sites based on oceanography and the sampling design described above.

```
make_environment <- function(n_sites = 10,
                             ocean_temp = 2,
                             temp_sd = 0,
                             ocean_recruitment = -2,
                             recruitment_sd = 0,
                             temp_mean = 15,
                             rec_mean = 10,
                             site_sd = 1,
                             seed = NULL){

  if(!is.null(seed)) set.seed(seed)

  tibble(
    site = as.character(1:n_sites)) %>%
    mutate(
      oceanography = rnorm(n_sites),
      site_temp = temp_mean +
        rnorm(n_sites, ocean_temp * oceanography, temp_sd),
      site_recruitment = rec_mean +
        rnorm(n_sites, ocean_recruitment * oceanography, recruitment_sd),
      site_int = rnorm(n_sites, 0, sd = site_sd)

    )
}
```

Great. Now, we need to add that year-to-year or plot-to-plot variability.

```
make_plots <- function(sites_df,
                       n_plots_per_site = 10,
                       plot_temp_sd = 1,
                       temp_effect = 1,
                       recruitment_effect = 1,
                       sd_plot = 2,
                       seed = NULL){

  if(!is.null(seed)) set.seed(seed)

  sites_df %>%
```

```
    rowwise() %>%
    mutate(
      plot_temp_dev_actual = list(rnorm(n_plots_per_site,
                                        0, plot_temp_sd))) %>%
    unnest(plot_temp_dev_actual) %>%
    mutate(
      plot_temp = site_temp + plot_temp_dev_actual,
      snails = rnorm(n(),
                     temp_effect*plot_temp +
                       recruitment_effect*site_recruitment +
                       site_int,
                     sd_plot)) %>%
    ungroup() %>%
    group_by(site) %>%
    mutate(year = 1:n(),
           site_mean_snails = mean(snails),
           site_mean_temp = mean(plot_temp),
           plot_snail_dev = snails - site_mean_snails,
           plot_temp_dev = plot_temp - site_mean_temp,
           site_mean_snail = mean(snails),
           site_snail_dev = snails - mean(snails),
           delta_snails = snails - lag(snails),
           delta_temp = plot_temp - lag(plot_temp)) %>%
    ungroup()

}
```

To analyze our data, we will compare several different fit models.

- A naive linear model with no site term
- A random effects using site as a random intercept
- A fixed effects model where site is a fixed effect (i.e., turned into 1/0 dummy variables)
- A model where we include the site mean temperature as a covariate and site is a random effect
- A model where we include site mean temperature as a covariate and site mean centered temperature (i.e., temperature at a site in a year minus it's mean over the entire data set). Site is included as a random effect
- A panel model where we look at change in snails between years versus change in temperature between years with a random site effect

```
analyze_plots <- function(plot_df){

  m <-  tribble(
```

```
    ~model_type, ~fit,
    "Naive", lm(snails ~ plot_temp, data = plot_df),
    "RE", lmer(snails ~ plot_temp + (1|site), data = plot_df),
    "FE Using Mean Differencing", lm(plot_snail_dev ~ plot_temp_dev, data =
plot_df), #fix SE?
    "FE with Dummy Variables", lm(snails ~ plot_temp + site, data = plot_df),
    "Group Mean Covariate", lmer(snails ~ plot_temp + site_mean_temp +
(1|site), data = plot_df),
    "Group Mean Centered", lmer(snails ~ plot_temp_dev + site_mean_temp +
(1|site), data = plot_df),
    "Group Mean Covariate, no RE", lm(snails ~ plot_temp + site_mean_temp,
data = plot_df),
    "Group Mean Centered, no RE", lm(snails ~ plot_temp_dev + site_mean_temp,
data = plot_df),
    "First Differences", lm(delta_snails ~ delta_temp,data = plot_df) #fix
SE?

  ) %>%
    mutate(coefs = map(fit, tidy), #get coefficients with broom
           out_stats = map(fit, glance),
           temp_effect = map(coefs, get_temp_coef),
           model_type = fct_inorder(model_type))

  m
}

get_temp_coef <- function(a_tidy_frame){
  a_tidy_frame %>%
    filter(term %in% c("plot_temp", "plot_temp_dev", "delta_temp")) %>%
    select(estimate, std.error)
}
```

#### Simulations and Results

Let's begin by setting up 100 replicate simulations.

```
set.seed(31415)
n_sims <- 100

envt <- tibble(
  sims = 1:n_sims
) %>%
  mutate(sites = map(sims, ~make_environment()))
```

Just for a sanity check, here's the relationship between temperature and recruitment at the site level across all simulations.

```
ggplot(envt %>%
         unnest(sites),
       aes(x = site_temp, y = site_recruitment, group = sims)) +
```

```r
  geom_point(alpha = 0.4, color = 'grey') +
  stat_smooth(method = "lm", size = 0.5, alpha = 0.5,
              fill = NA, color = "black") +
  labs(x = "Site Average Temperature",
       y = "Site Recruitment of Individuals Per Plot")
```

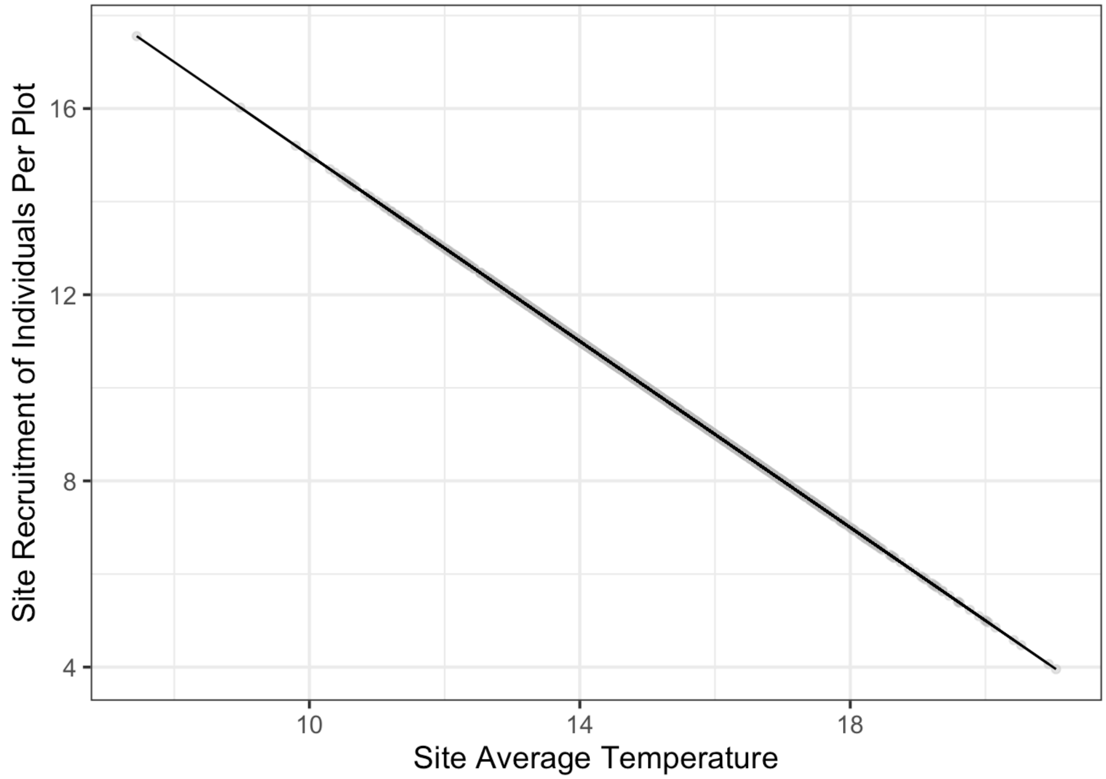


Great! Now, let's setup our sampling over time.

```r
plots_df <- envt %>%
  mutate(site_year = map(sites, make_plots))
```

And, again, a sanity check…

```r
ggplot(plots_df %>%
         unnest(site_year),
       aes(x = plot_temp, y = snails, color = as.character(sims), group =
sims)) +
  stat_smooth(method = "lm", fill = "NA") +
  guides(color = "none") +
  labs(x = "Plot Temperature",
       y = "Snails per Plot")
```

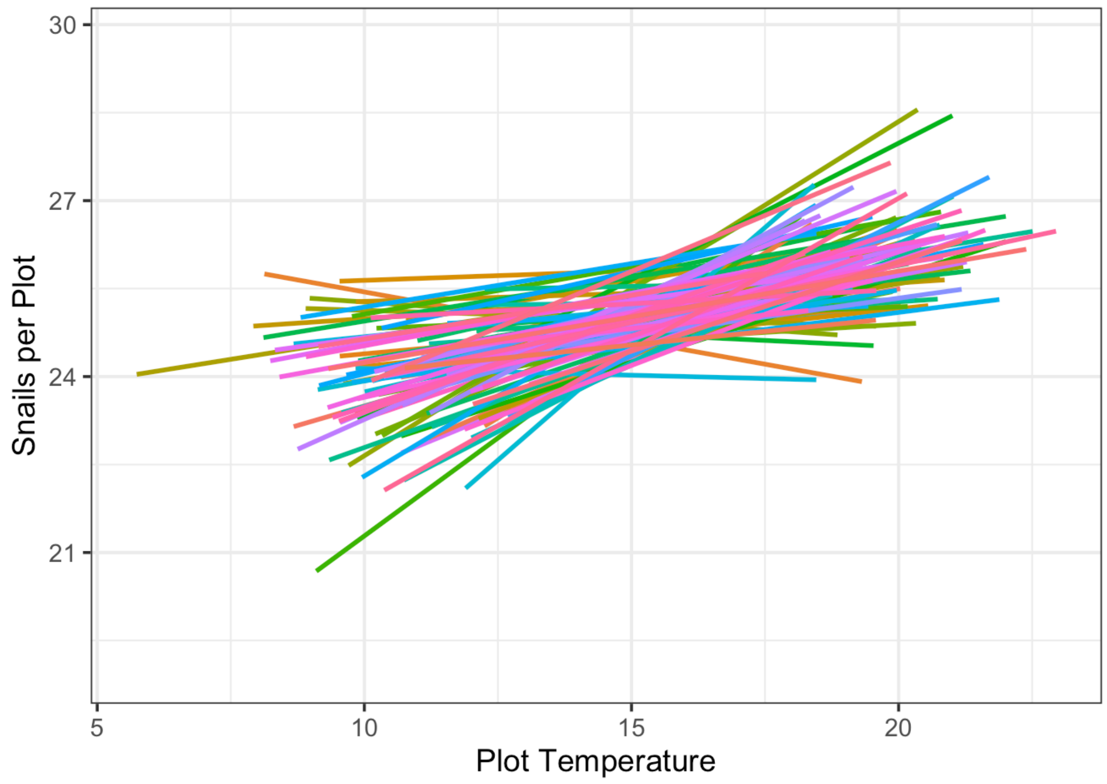


So we can see that in this setup, the snail-temperature relationship is positive in nearly all of the simulations. But, how positive is it? What would our coefficients show on average across simulations?

Let's fit models to each set of data

```
analysis_df <- plots_df %>%
  mutate(analysis = map(site_year, analyze_plots)) %>%
  unnest(analysis)
```

And now let's look at the distribution of coefficients that would describe the relationship between temperature and snails from each mode.

```
analysis_df %>%
  unnest(temp_effect) %>%
  ggplot(aes(y = fct_rev(model_type), x = estimate)) +
  ggridges::stat_density_ridges() +
  labs(y="", x = "Model Estimated Temperature Effect") +
  geom_vline(xintercept = 1, linewidth = 1.5, lty = 3)
```

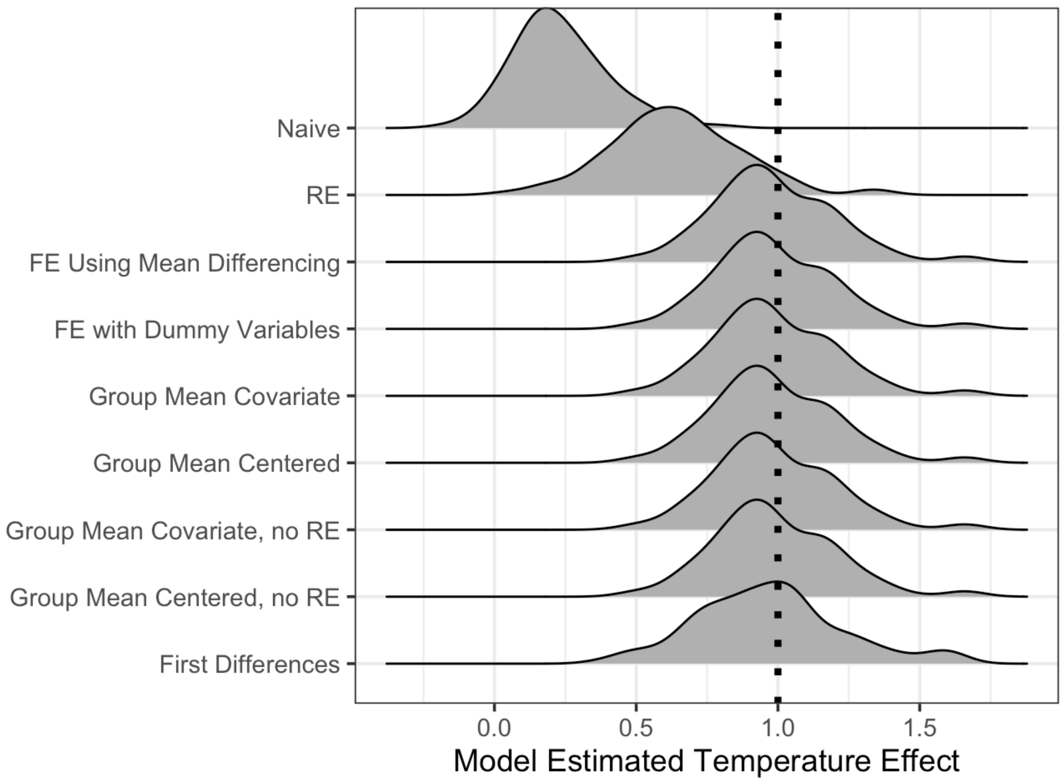


Eyeballing it, we can see of course the naive model is too low, as is the RE model. How bad is the bias for the RE model?

```
analysis_df %>%
  unnest(temp_effect) %>%
  group_by(`Model Type` = model_type) %>%
  summarize(`Mean Estimate` = mean(estimate),
            `SD Estimate` = sd(estimate)) %>%
  knit_table
```

| Model Type | Mean Estimate | SD Estimate |
|---|---|---|
| Naive | 0.231 | 0.165 |
| RE | 0.640 | 0.232 |
| FE Using Mean Differencing | 0.985 | 0.215 |
| FE with Dummy Variables | 0.985 | 0.215 |
| Group Mean Covariate | 0.985 | 0.215 |
| Group Mean Centered | 0.985 | 0.215 |
| Group Mean Covariate, no RE | 0.985 | 0.215 |
| Group Mean Centered, no RE | 0.985 | 0.215 |
| First Differences | 0.971 | 0.259 |

The downward bias produced by poor model choice is clear. We can see it if we plot the coefficient minus one, which if unbiased should reveal a distribution centered on 0.

```r
analysis_df %>%
  unnest(temp_effect) %>%
  ggplot(aes(y = fct_rev(model_type), x = estimate - 1)) +
  ggridges::stat_density_ridges() +
  labs(y="", x = "Bias")
```

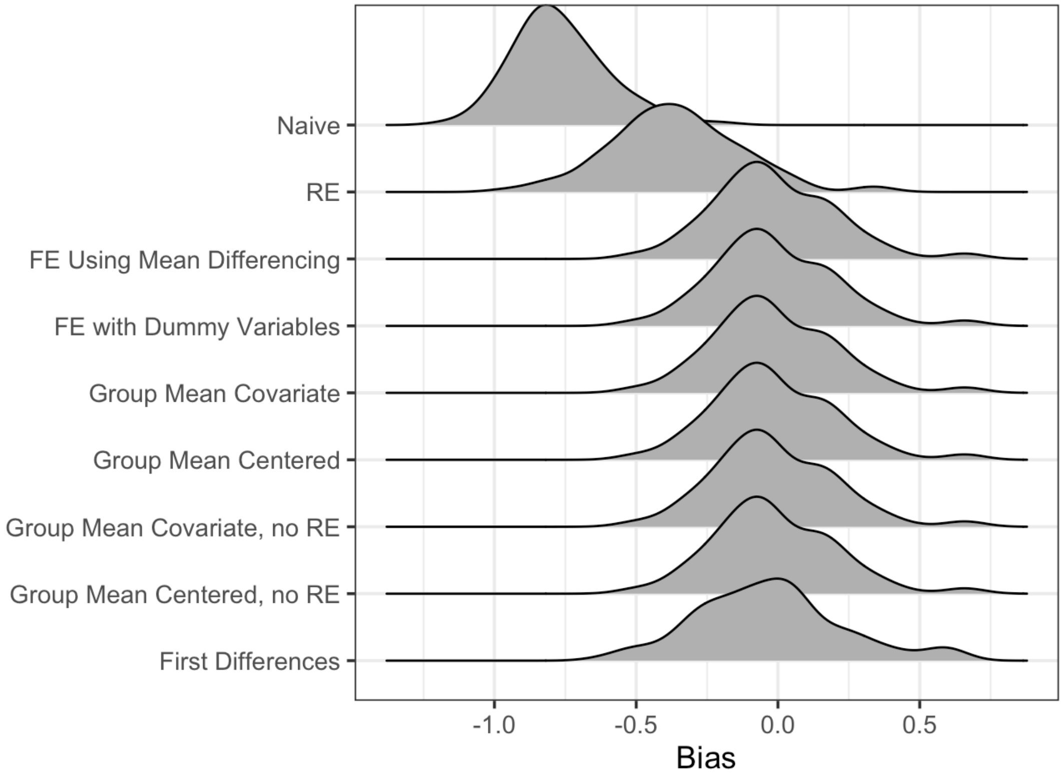


We can also look at the distribution of, for each simulated data set, how different the RE model is from each other model.

```r
analysis_df %>%
  filter(model_type != "Naive") %>%
  unnest(temp_effect) %>%
  group_by(sims) %>%
  mutate(diff_from_re = estimate - estimate[1]) %>%
  ungroup() %>%
  filter(model_type != "RE") %>%
  ggplot(aes(y = fct_rev(model_type), x = diff_from_re)) +
  ggridges::stat_density_ridges() +
  labs(y="", x = "Difference from RE Model\nTemperature Effect")
```

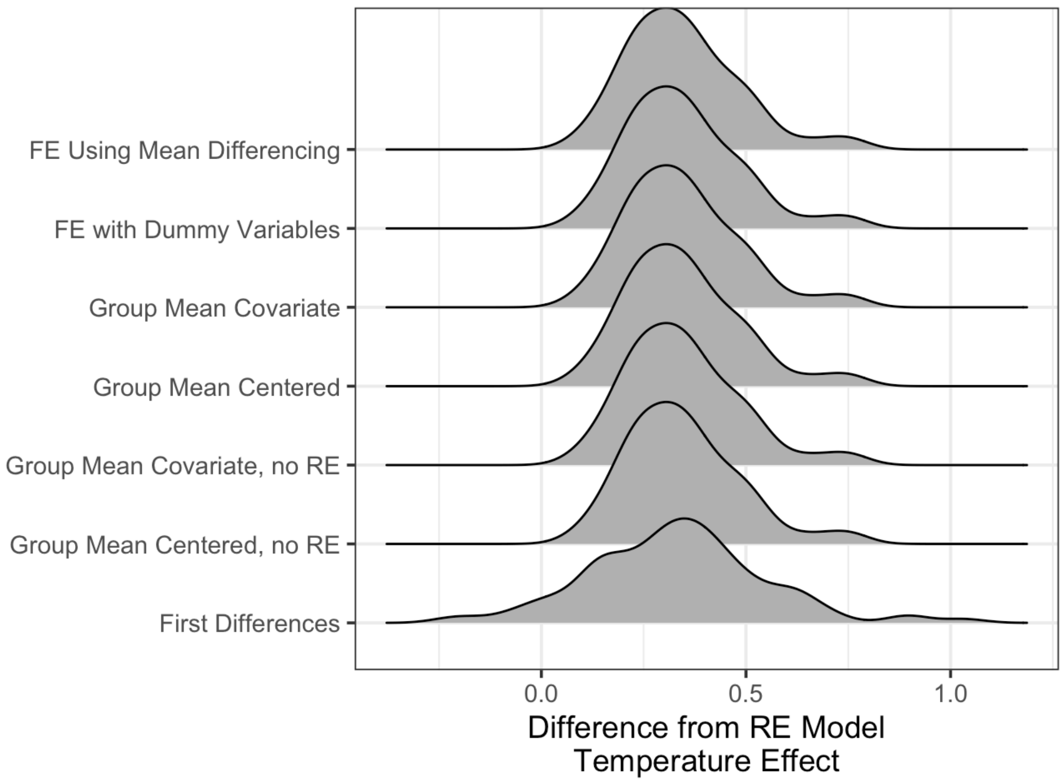


Note, in all cases, we can see the effects of downward bias. This is clear, but, to put it in numbers -

```
analysis_df %>%
  filter(model_type != "Naive") %>%
  unnest(temp_effect) %>%
  group_by(sims) %>%
  mutate(diff_from_re = estimate - estimate[1]) %>%
  ungroup() %>%
  filter(model_type != "RE") %>%
  group_by(model_type) %>%
  summarize(`Mean Diff from RE` = mean(diff_from_re),
            `SD Diff from RE` = sd(diff_from_re))  %>%
  knit_table
```

| model_type | Mean Diff from RE | SD Diff from RE |
|---|---|---|
| FE Using Mean Differencing | 0.346 | 0.138 |
| FE with Dummy Variables | 0.346 | 0.138 |
| Group Mean Covariate | 0.346 | 0.138 |
| Group Mean Centered | 0.346 | 0.138 |
| Group Mean Covariate, no RE | 0.346 | 0.138 |
| Group Mean Centered, no RE | 0.346 | 0.138 |
| First Differences | 0.331 | 0.218 |

If we were doing straight hypothesis testing, how often would our estimate of the temperature coefficient either overlap 0 or not have 1 within its confidence interval?

```
analysis_df %>%
  filter(model_type != "Naive") %>%
  unnest(temp_effect) %>%
  mutate(overlap_0 = (estimate - 2*std.error)<0,
         overlap_1 = (estimate - 2*std.error)<1 &
           (estimate + 2*std.error)>1) %>%
  group_by(`Model Type` = model_type) %>%
  summarize(`95% CI Contains 0` = sum(overlap_0)/n(),
            `95% CI does Not Contain 1` = (n()-sum(overlap_1))/n()) %>%
  knit_table
```

| Model Type | 95% CI Contains 0 | 95% CI does Not Contain 1 |
|---|---|---|
| RE | 0.08 | 0.54 |
| FE Using Mean Differencing | 0.00 | 0.05 |
| FE with Dummy Variables | 0.00 | 0.05 |
| Group Mean Covariate | 0.00 | 0.05 |
| Group Mean Centered | 0.00 | 0.05 |
| Group Mean Covariate, no RE | 0.01 | 0.04 |
| Group Mean Centered, no RE | 0.01 | 0.04 |
| First Differences | 0.01 | 0.12 |

Here we see the RE model is more likely to be subject to type II error. Further, it is far more likely than any other technique to not have the true coefficient value within 2 CI of its estimand.

#### A Wrapper Function for Simulation

This has been useful, but, if we want to automate the process for further exploration, let's wrap the code above into a function.

```
make_sims_and_analyze <- function(n_sims = 100,
                                  n_sites = 10,
                                  ocean_temp = 2,
                                  temp_sd = 0,
                                  ocean_recruitment = -2,
                                  recruitment_sd = 0,
                                  temp_mean = 15,
                                  rec_mean = 10,
                                  site_sd = 1,
                                  n_plots_per_site = 10,
                                  plot_temp_sd = 1,
                                  temp_effect = 1,
                                  recruitment_effect = 1,
                                  sd_plot = 2,
```

```
                                seed = NULL) {
  #should we set a seed?
  if (!is.null(seed))
    set.seed(seed)

  # make an envt data frame
  out_df <- tibble(sims = 1:n_sims) %>%
    mutate(
      sites = map(
        sims,
        ~make_environment(
         n_sites = n_sites,
          ocean_temp = ocean_temp,
          temp_sd = temp_sd,
          ocean_recruitment = ocean_recruitment,
          recruitment_sd = recruitment_sd,
          temp_mean = temp_mean,
          rec_mean = rec_mean,
          site_sd = site_sd)
      )
    ) %>%
    #now add plots
    mutate(
      site_year = map(
        sites,
        make_plots,
        n_plots_per_site = n_plots_per_site,
        plot_temp_sd = plot_temp_sd,
        temp_effect = temp_effect,
        recruitment_effect = recruitment_effect,
        sd_plot = sd_plot
      )
    ) %>%

    #and analysis
    mutate(analysis = map(site_year, analyze_plots)) %>%
    unnest(analysis)

}
```

#### Is that Random Effect Needed?

If we look at the correlated random effects models, what's the RE?

```
analysis_df %>%
  filter(model_type %in% c("Group Mean Covariate", "Group Mean Centered"))
%>%
  unnest(coefs) %>%
  filter(group == "site") %>%
  group_by(`Model Type` = model_type) %>%
```

```
  summarize(Term = "Site Random Effect",
            `Mean Site SD` = mean(estimate),
            `SD in Site SD` = sd(estimate)) %>%
  knit_table
```

| Model Type | Term | Mean Site SD | SD in Site SD |
|---|---|---|---|
| Group Mean Covariate | Site Random Effect | 0.977 | 0.438 |
| Group Mean Centered | Site Random Effect | 0.977 | 0.438 |

Both are the same, which makes sense given the formulation of the model. What if there was no additional site-level variation uncorrelated with temperature, though?

```
re_0_frame <- make_sims_and_analyze(site_sd = 0)

re_0_frame %>%
  filter(model_type %in% c("Group Mean Covariate", "Group Mean Centered"))
%>%
  unnest(coefs) %>%
  filter(group == "site") %>%
  group_by(`Model Type` = model_type) %>%
  summarize(Term = "Site Random Effect",
            `Mean Site SD` = mean(estimate),
            `SD in Site SD` = sd(estimate)) %>%
  knit_table
```

| Model Type | Term | Mean Site SD | SD in Site SD |
|---|---|---|---|
| Group Mean Covariate | Site Random Effect | 0.273 | 0.261 |
| Group Mean Centered | Site Random Effect | 0.273 | 0.261 |

Both REs are the same - again - but both overlap 0. In the system we simulated, there is no uncorrelated site-level variability. It's all at the site-year (or site-plot) level. We have indeed run these models with *no site random effect*. They produce the same answers for coefficients - both in this simulation with no site-level random effect as well as above when there *was* a site-level random effect.

```
# Without random site variation
re_0_frame %>%
  filter(grepl("Group Mean", model_type)) %>%
  unnest(temp_effect) %>%
  group_by(`Model Type` = model_type) %>%
  summarize(`Mean Temp Effect` = mean(estimate),
            `SD Temp Effect` = sd(estimate)) %>%
  knit_table
```

| Model Type | Mean Temp Effect | SD Temp Effect |
|---|---|---|
| Group Mean Covariate | 0.969 | 0.203 |
| Group Mean Centered | 0.969 | 0.203 |

| Model Type | Mean Temp Effect | SD Temp Effect |
| --- | --- | --- |
| Group Mean Covariate, no RE | 0.969 | 0.203 |
| Group Mean Centered, no RE | 0.969 | 0.203 |

```
# With random site variation
analysis_df %>%
  filter(grepl("Group Mean", model_type)) %>%
  unnest(temp_effect) %>%
  group_by(`Model Type` = model_type) %>%
  summarize(`Mean Temp Effect` = mean(estimate),
            `SD Temp Effect` = sd(estimate)) %>%
  knit_table
```

| Model Type | Mean Temp Effect | SD Temp Effect |
| --- | --- | --- |
| Group Mean Covariate | 0.985 | 0.215 |
| Group Mean Centered | 0.985 | 0.215 |
| Group Mean Covariate, no RE | 0.985 | 0.215 |
| Group Mean Centered, no RE | 0.985 | 0.215 |

So should we just not worry about a site-level random effect? Not necesarily. First, if we consider a site RE, we can look at residual variation due to both between site differences as well as residual replicate-level variation. This can be a useful analysis when attempting to tease apart variation that is versus is not correlated with a driver of interest. Second, while models without a site random effect can be fit and used, these models will be structurally incorrect - replicates are not IID, as there is correlated error within sites. Last, mixed models can also provide advantages when handling models with unbalanced data between sites (see below).

Practically, however, the difference comes in with respect to whether you are interested in between site variation or not. The residual inflates with no RE, as it is now the combination of between site residual and within site residual terms. This could make a difference for various statistical tests down the line as well, but in terms of parameter estimates, we are still estimating a clean causal effect.

#### What About Unbalanced Data?

One of the advantages to mixed modeling approaches is how they handle unbalanced data. What if, in the above example, we had lost samples from each site generating unbalanced data?

```
set.seed(31415)
#what do we keep?
years_to_keep <- round(runif(10, 3, 10)) %>%
    imap_dfr( ~ tibble(site = as.character(.y),
                       year = sample(1:10, .x),
                       keep = T))

#a function for unbalancing
```

```r
unbalance <- function(site_year_df, keep_df){
  left_join(site_year_df, keep_df) %>%
    filter(keep)
}

#apply keeping to each site-year
unbalanced_df <- analysis_df %>%
  select(-model_type, -fit, -coefs, -out_stats, -temp_effect) %>%
  mutate(site_year = map(site_year, unbalance,
                              keep_df = years_to_keep),
         analysis = map(site_year, analyze_plots)) %>%
    unnest(analysis)
```

How does this lack of balance affect estimation of causal coefficients?

```r
unbalanced_df %>%
    unnest(temp_effect) %>%
  group_by(`Model Type` = model_type) %>%
  summarize(`Mean Estimate` = mean(estimate),
            `SD Estimate` = sd(estimate)) %>%
  knit_table
```

| Model Type | Mean Estimate | SD Estimate |
|---|---|---|
| Naive | 0.245 | 0.195 |
| RE | 0.500 | 0.297 |
| FE Using Mean Differencing | 0.986 | 0.275 |
| FE with Dummy Variables | 0.990 | 0.281 |
| Group Mean Covariate | 0.986 | 0.281 |
| Group Mean Centered | 0.986 | 0.281 |
| Group Mean Covariate, no RE | 0.982 | 0.299 |
| Group Mean Centered, no RE | 0.982 | 0.299 |
| First Differences | 0.961 | 0.316 |

We can see that point estimation here is improved somewhat. Although this could vary. More telling would be an exploration of those group means in the mixed versus fixed models.
