## Supplemental Tables, Figures, Text, Data, and Code for "Causal inference with observational data and unobserved confounding variables": supporting_information_s7.pdf

### Supporting Information S7: Implementing OVB Model Methods in R

#### Introduction

To demonstrate the simplicity of the approaches outlined in the manuscript, here we present an analysis of a single simulated data set that stems from the simulations in the text. We have provided a data file (Supplementary Data 1) for ease of use[1]. To start, we will load a few libraries. If you do not have them installed already, you can do so via `install.packages()` or substitute `pacman::p_load()` for `library()` below after installing `pacman`.

For those not familiar with the packages used in this example, here we describe what purpose each one serves. We will use `dplyr` for making sure columns are of the right class and for the action of calculating group meand and anomalies. `purrr` will be used to automate some summary output creation later in the script. `lme4` will be used to fit linear mixed effects models. Last, `broom.mixed` is a wonderful package that generates easily readable output from mixed models.

```
## Libraries
# data manipulation
library(dplyr) # used version 1.1.4
library(purrr) # used version 1.0.2

# modeling
library(lme4) # used version 1.1-35.1
library(fixest) # version 0.11.2

# viewing output
library(broom) # used version 1.0.5
library(broom.mixed) # used version 0.2.9.4
```

First, we will load the data and turn the site variable into a character. We note that it is common to forget to make items like 'site' into categorical variables, and this can have disastrous consequences. We then output the first six lines of the data to show what it looks like. Interested readers are welcome to explore using other methods (we are fans of the `visdat` and `skimr` packages as well as just using `str()`).

```
dat <- read.csv("data/sim_snails_seed_ovb.csv") |>
  mutate(site = as.character(site))

head(dat)

  site year     temp   snails
1    1    1 13.68623 25.62675
2    1    2 15.45278 23.87720
```

```
3    1    3 13.67513 23.96380
4    1    4 15.09723 27.00633
5    1    5 15.13115 28.56318
6    1    6 15.11702 31.11580
```

#### Models

With the data as shown above, we can now fit a naive linear model that does not incorporate site using `lm()`.

```r
mod_naive <- lm(snails ~ temp, data = dat)
```

We can then fit an Random Effects form of the model with site as the random effect (RE) using `lmer()` from `lme4`.

```r
mod_re <- lmer(snails ~ temp + (1|site),
               data = dat)
```

To fit an Econometric Fixed Effects model, we can again use `lm()` and incorporate site as a Fixed Effect predictor. First, make sure that site is a character of factor, though, as we did above when we loaded the data. If you are unsure, check the class of the site column.

```r
class(dat$site) # all good!
```

```
[1] "character"
```

```r
mod_fe <- lm(snails ~ temp + site, data = dat)
```

To see the dummy coding of the FE, try `model.matrix(mod_fe)`. To implement a two-way fixed effects model, you can add site + year to the formula above, making sure year is also a character or factor.

We can implement a FE model using the Fixed Effects Transformation as well. To do this, we group by site and then calculate the site-level anomaly for both snails and temperature. We then fit the linear model with `lm()` examining the relationship between these two site-level anomalies.

```r
dat <- dat |>
  group_by(site) |>
  mutate(snail_site_anom = snails - mean(snails),
         temp_site_anom = temp - mean(temp)) |>
  ungroup()

# FE Transformation Model
mod_fe_trans <- lm(snail_site_anom ~ temp_site_anom, data = dat)
```

Econometric Fixed Effects can also be implemented in the package fixest with the function `feols()`, which is highly efficient for large datasets with many units (e.g. sites) and therefore lots of fixed effects. It also easily incorporates clustered robust standard errors and other standard errors. In fact, it uses clustered SE by default. Here, we force it to produce non-clustered SEs for comparison to other approaches. See our section on robust standard errors below for more.

```
mod_fe_feols <- feols(snails ~ temp | site, se = "standard", data = dat)
```

Like the FE models above, you can have a TWFE model by having the FE specification as `site + year`.

For the Group Mean Covariate (Mundlak) model and the Group Mean Centered model, we need to calculate a mean temperature by site. Then we can fit both models with that mean as a hierarchical predictor and a random effect of site using `lmer()`.

```
dat <- dat |>
  group_by(site) |>
  mutate(site_mean_temp = mean(temp)) |>
  ungroup()

# Group Mean Covariate Model
mod_gmcov <- lmer(snails ~ temp + site_mean_temp + (1|site),
                  data = dat)

# Group Mean Centered Model
mod_gmcent <- lmer(snails ~ temp_site_anom + site_mean_temp + (1|site),
                  data = dat)
```

#### Comparison

Let's compare the performance of these different models as estimators of the temperature effect. Here, we create a named list of models, and then use `purrr::map()` functions along with `tidy()` from `broom` and `broom.mixed` to get well formatted output.

```
mods <- list(mod_naive = mod_naive,
             mod_re = mod_re,
             mod_fe = mod_fe,
             mod_fe_trans = mod_fe_trans,
             mod_fe_feols = mod_fe_feols,
             mod_gmcov = mod_gmcov,
             mod_gmcent = mod_gmcent)

out <- map(mods, tidy) |>
  map_dfr(~ .x |> select(term, estimate, std.error),
          .id = "model") |>
  filter(term %in% c("temp", "temp_site_anom"))

out
```

```
# A tibble: 7 × 4
  model         term            estimate std.error
  <chr>         <chr>              <dbl>     <dbl>
1 mod_naive     temp               0.164     0.105
```

```
2 mod_re       temp               0.579     0.169
3 mod_fe       temp               0.955     0.213
4 mod_fe_trans temp_site_anom     0.955     0.203
5 mod_fe_feols temp               0.955     0.213
6 mod_gmcov    temp               0.955     0.213
7 mod_gmcent   temp_site_anom     0.955     0.213
```

#### Robust Standard Errors

Robust standard errors are of use in a great many cases. If errors are hetergeneous between groups, have temporal autocorrelation, or other issues, they can be very used Oshchepkov and Shirokanova (2022). In R, two main ways to use them are either via the `sandwich` package along with `lmtest` or to fit them with the `fixest` package.

```
library(lmtest) # version 0.9-40
library(sandwich) # version 3.1-0
library(fixest) # version 0.11.2
```

We can look at the SE for the temperature effect from `mod_fe` first with no correction, then the Huber-White correction from `sandwich`, and finally we can fit the same model using `fixest::feols()`, which uses slightly different syntax for incorporating fixed effects, and also output the Huber-White correction. See also https://cran.r-project.org/web/packages/fixest/vignettes/standard_errors.html for more on SE in `fixest`.

Note, there is a small difference between the output from `sandwich` and `fixest` for clustering and the Huber-White correction together due to how each package implements the calculation of effective degrees of freedom.

```
# No correction
tidy(mod_fe) |>
  filter(term == "temp")
```

```
# A tibble: 1 × 5
  term   estimate std.error statistic   p.value
  <chr>     <dbl>     <dbl>     <dbl>     <dbl>
1 temp      0.955     0.213      4.49 0.0000215
```

```
# Huber-White SE without clustering
coeftest(mod_fe,
         vcov = vcovCL(mod_fe,
                       type = "HC1")) |>
    tidy() |>
    filter(term == "temp")
```

```
# A tibble: 1 × 5
  term   estimate std.error statistic    p.value
  <chr>     <dbl>     <dbl>     <dbl>      <dbl>
1 temp      0.955     0.197      4.85 0.00000522
```

```
# Using fixest for Huber-White correction without clustering
feols(snails ~ temp | site,
      vcov = "hetero",
      data = dat) |>
  tidy()

# A tibble: 1 × 5
  term  estimate std.error statistic    p.value
  <chr>    <dbl>     <dbl>     <dbl>      <dbl>
1 temp     0.955     0.197      4.85 0.00000522

# Huber-White SE with site-level clustering via the Sandwich package in R
coeftest(mod_fe,
         vcov = vcovCL(mod_fe,
                       cluster = ~ site,
                       type = "HC1")) |>
  tidy() |>
  filter(term == "temp")

# A tibble: 1 × 5
  term  estimate std.error statistic       p.value
  <chr>    <dbl>     <dbl>     <dbl>         <dbl>
1 temp     0.955     0.143      6.66 0.00000000222

# Using fixest for Huber-White correction and site-level clustering with
feols
feols(snails ~ temp | site,
      vcov = "cluster",
      data = dat) |>
  tidy(vcov = "cluster")

# A tibble: 1 × 5
  term  estimate std.error statistic   p.value
  <chr>    <dbl>     <dbl>     <dbl>     <dbl>
1 temp     0.955     0.137      6.99 0.0000643
```

#### Footnotes

1. Of note, we generated it using the functions `make_environment()` piped to `make_plots()` in Appendix A. We set a seed to make a reproducible example. The number for the seed was generated by `TeachingDemos::char2seed("OVB")` which translated to 834.↩
